# Quantitative RNA pseudouridine landscape reveals dynamic modification patterns and evolutionary conservation across bacterial species

**DOI:** 10.1101/2025.05.13.653745

**Authors:** Letong Xu, Shenghai Shen, Yizhou Zhang, Zhihao Guo, Beifang Lu, Jiadai Huang, Runsheng Li, Yitong Shen, Li-Sheng Zhang, Xin Deng

**Author notes:** Correspondence (Xin Deng) and (Li-Sheng Zhang). These authors contributed equally to the paper as the first authors.

## Abstract

Pseudouridine (Ψ) modifications are the most abundant RNA modifications; however, their distribution and functional significance in bacteria remain largely unexplored compared to eukaryotic systems. In this study, we present the first transcriptome-wide and quantitative mapping of Ψ modifications across five diverse bacterial species (*Bacillus cereus, Escherichia coli, Klebsiella pneumoniae, Pseudomonas aeruginosa*, and *Pseudomonas syringae*) at single-base resolution, utilizing the optimized BID-seq method for bacterial RNA. Our analysis revealed growth phase-dependent dynamics of pseudouridylation in bacterial tRNA and mRNA, particularly in genes enriched in core metabolic pathways. Comparative analysis demonstrated evolutionarily conserved features of Ψ modifications, such as dominant motif contexts, Ψ clustering within operons, etc. Functional analysis indicated Ψ modifications influence bacterial mRNA stability, translation, and interactions with specific RNA-binding proteins (RBPs) in response to changing cellular demands during growth phase transitions. The integrated computational analysis on local RNA architecture was conducted to elucidate the structure-dependent Ψ modifications in bacterial RNA. Furthermore, we developed an integrated deep learning framework, combining Transformer-GNN-based neural networks (pseU_NN) to capture both RNA sequence and structural features for effective prediction of Ψ-modified sites. Overall, our study provides valuable insights into the landscapes of bacterial RNA Ψ modifications and establishes a foundation for future mechanistic investigations into the functions of Ψ in bacterial RNA regulation.

## Introduction

RNA modifications are a crucial layer of post-transcriptional regulation in biological systems, with approximately 170 distinct chemical modifications identified to date(Roundtree et al., 2017). Pseudouridine (Ψ), often referred to as the “fifth nucleoside,” is one of the most prevalent and evolutionarily conserved RNA modifications(Cerneckis et al., 2022; Rodell et al., 2024). This unique modification arises from a specific isomerization process in which uridine undergoes a site-specific intramolecular rearrangement(Yu and Allen, 1959). Ψ can thermodynamically stabilize RNA structures by enhancing base stacking and increasing the rigidity of the sugar-phosphate backbone, which helps maintain the structural folding of functional RNAs such as tRNA and rRNA(Pan et al., 2003; Roovers et al., 2006). For example, Ψ at the 55^th^ position of tRNA, a universally conserved modification in both eukaryotes and prokaryotes, is crucial for regulating tRNA stability and aminoacylation levels(Ishida et al., 2011; Schultz et al., 2024). Ψ modifications in rRNA play important roles in rRNA biogenesis and function, as well as in mRNA translation in both mammalian cells and bacteria(Leppik et al., 2017; Sloan et al., 2017; Zhao et al., 2023).

The regulation of mRNA translation is highly complex, and mRNA stability significantly influences gene expression in humans. Ψ-modified mRNAs demonstrate enhanced stability due to their resistance to RNase L-mediated degradation(Anderson et al., 2011). Pre-mRNA is found to be pseudouridylated co-transcriptionally, with Ψ enriched near alternative splicing regions and RNA-binding protein (RBP) binding sites(Martinez et al., 2022). Moreover, Ψ located within exon regions can alter codon properties to modulate translation, while Ψ modifications at stop codons promote ribosomal readthrough(Hoernes et al., 2016; Karijolich and Yu, 2012, 2011). Ψ also facilitates the low-level synthesis of peptide products from individual mRNA sequences in human cells and increases the rate at which near-cognate tRNA^Val^ interacts with ΨUU codons(Eyler et al., 2019), suggesting a more complex regulatory role for Ψ in translation.

Until recently, comprehensive investigations of bacterial RNA pseudouridylation have been limited due to technical challenges in precisely mapping and quantifying Ψ modifications at single-nucleotide resolution. While eukaryotic mRNA can be easily isolated and assessed using poly-A selection methods, bacterial transcripts lack poly-A tails and are predominantly composed of ribosomal RNA, which accounts for over 95% of total RNA(Liang et al., 2000). This methodological gap has hindered the functional exploration of Ψ roles in bacterial RNA. A previously reported CMC-based method can selectively label Ψ sites and produce truncation signatures during reverse transcription. However, this method often exhibits several drawbacks, including relatively low sensitivity and limitations in quantifying Ψ modification levels. Recently, a new technique named Bisulfite-induced deletion sequencing (BID-seq) has emerged, utilizing unique deletion signatures induced at Ψ-modifed sites to achieve base-resolution and quantitative characterization of Ψ sites across the transcriptome(Dai et al., 2023; Zhang et al., 2024).

To address these challenges and conduct an in-depth study of bacterial Ψ modifications, we developed an optimized BID-seq method for bacterial RNA, termed baBID-seq. We selected *Klebsiella pneumoniae* (*K. pneumoniae), Bacillus cereus* (*B. cereus), Pseudomonas aeruginosa* (*P. aeruginosa)*, and *Pseudomonas syringae* (*P. syringae)* based on their biological relevance and taxonomic diversity. *K. pneumoniae, B. cereus*, and *P. aeruginosa* are clinically important human pathogens responsible for a broad spectrum of infectious diseases, yet transcriptome-wide pseudouridylation has not been systematically characterized in these organisms(Ehling-Schulz et al., 2019; Kerr and Snelling, 2009; Wyres et al., 2020). *P. syringae*, a well-studied plant pathogen, was included to extend the analysis beyond human pathogens and to explore pseudouridine modification in a distinct ecological context(Xin et al., 2018). Collectively, these species encompass both Gram-positive (B. cereus) and Gram-negative (*K. pneumoniae, P. aeruginosa*, and *P. syringae*) bacteria and exhibit substantial differences in genome size, GC content, and pathogenic lifestyle. This selection provides a comparative framework for investigating conserved and species-specific features of bacterial pseudouridylation across diverse lineages. By combining efficient rRNA depletion in baBID-seq, we expanded our quantitative Ψ analysis to these four representative bacterial species, examining both exponential and stationary growth phases. Our comprehensive analysis revealed the landscape of Ψ modifications on bacterial rRNA, tRNA, and mRNA, highlighting evolutionarily conserved Ψ features across bacterial strains. We investigated the sequence and structural properties of local mRNA regions that influence Ψ deposition. Dynamic Ψ modifications at specific sites in tRNA and mRNA were observed, showing distinct accumulation patterns during the stationary growth phase. In *P. syringae*, we explored the roles of Ψ under nutrient-deficient conditions and found a positive correlation between mRNA translation efficiency (TE) and Ψ intensity(Hua et al., 2024). In *P. aeruginosa*, we provided evidence for the potential regulatory functions of Ψ in promoting mRNA interactions with the Hfq chaperone (Trouillon et al., 2022). Furthermore, we employed a hybrid LSTM-attention-based graph neural network (GNN) classification approach, integrating RNA sequence and local structural features to predict potential Ψ modification sites. Collectively, our systematic analysis revealed a novel and dynamic landscape of Ψ modifications, uncovering evolutionarily conserved modification features and key motifs and structural elements within mRNA regions that impact Ψ modification.

## Results

### Optimized BID-seq quantitatively maps Ψ modification in bacterial rRNA and tRNA

To investigate Ψ modifications in bacteria, we primarily applied the standard BID-seq protocol(Zhang et al., 2024) to total RNA isolated from *E. coli* and *P. aeruginosa* during exponential and stationary growth phases. Ψ sites on rRNA were identified through deletion signatures at single-base resolution, and the observed deletion ratios were utilized to assess Ψ modification fraction. By characterizing Ψ sites with significantly higher deletion ratios in the ‘BID-seq treated’ samples compared to the ‘input’ samples, we detected 9 out of 10 known Ψ sites, as well as one additional Ψ site, that were significantly modified in 23S and 16S rRNA, respectively (**Fig. 1a,b**). For example, Ψ781 in 16S rRNA of *P. aeruginosa* exhibited a distinct deletion signature in the ‘BID-seq treated’ samples (**Supplementary Fig. 1c**). Although Ψ is known to play a critical role in regulating rRNA local structure and biogenesis(Leppik et al., 2017), bacterial RNA BID-seq (baBID-seq) pointed out that the number of Ψ-modified sites is noticeably lower than the typical >100 Ψ sites found in mammalian rRNA(Dai et al., 2023). We then compared the rRNA Ψ fraction in *E. coli* and *P. aeruginosa* across the two growth stages, and almost all Ψ sites are highly conserved, with only slight variations in Ψ site deletion ratio (**Fig. 1a,b**).

**Figure 1.**
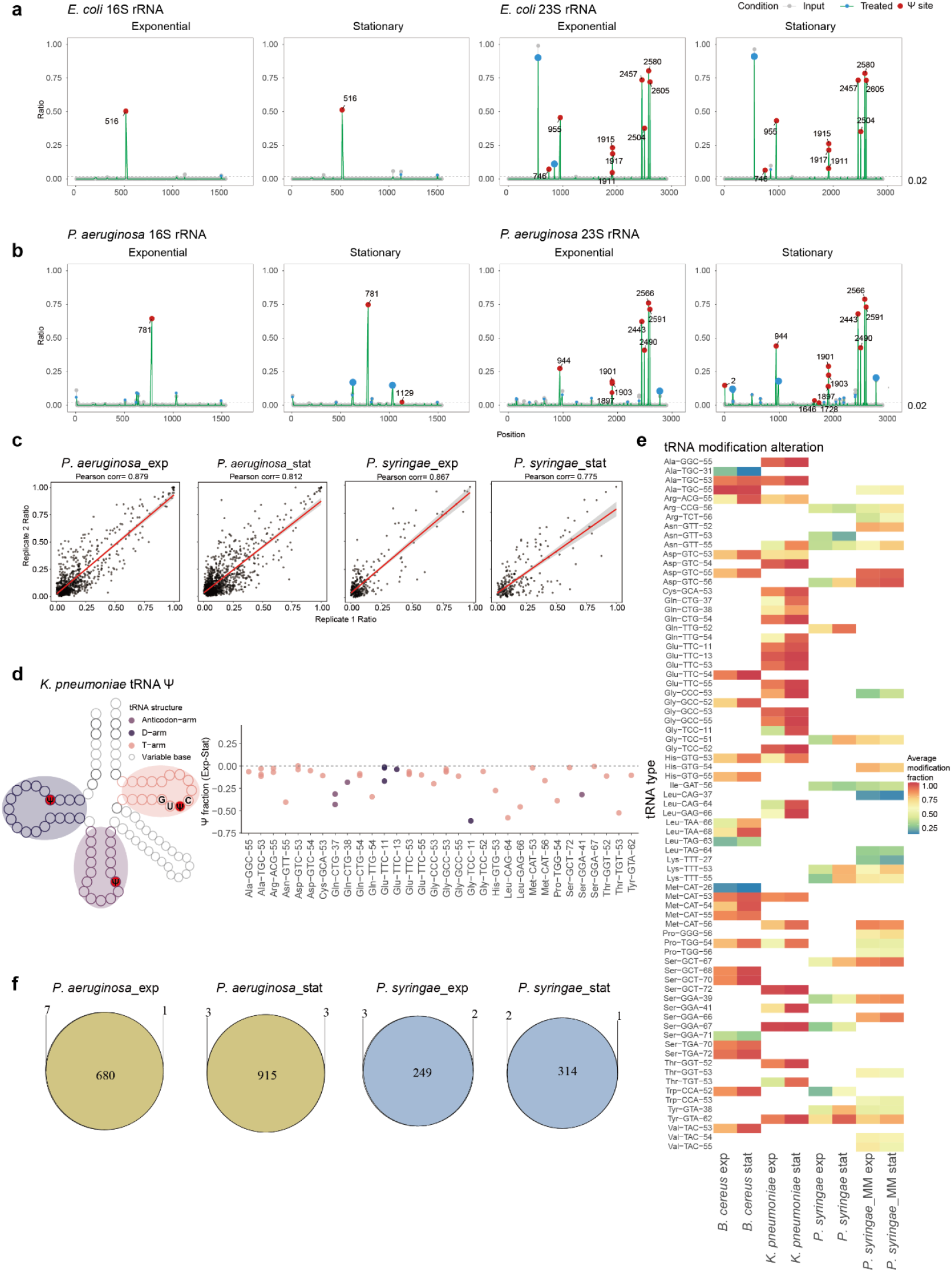
BID-seq identifies precise pseudouridine (Ψ) modification sites in ribosomal RNA and reveals dynamic Ψ modification patterns in transfer RNA. **a**,**b** Ψ modifications detected on 16S and 23S ribosomal RNA with baBID-seq of *E. coli* (**a**) and *P. aeruginosa* (**b**) total RNA during exponential and stationary growth phases. All Ψ sites in panels **a** and **b** were identified using filtration criteria of deletion fraction > 0.02 and *p*-value <= 1 × 10−^4^ **c** Pearson correlation analysis of Ψ modification fractions at individual sites between biological replicates. **d** Ψ modification fraction alteration pattern observed on specific sites in *K. pneumoniae* different tRNA regions. The left panel depicts the general tRNA secondary structure in *K. pneumoniae*. Structural regions are indicated in the legend and highlighted with corresponding colors. The right scatter plot’s y-axis depicts the Ψ fraction difference, calculated as (exponential phase Ψ fraction) - (stationary phase Ψ fraction). **e** Heatmap displaying the tRNA Ψ fraction alteration features detected across different conditions. Color represents the average Ψ fraction values (range from 2% to 100%) at specific sites within tRNA isoacceptors for each strain. The tRNA tags (labeled in y-axis) comprise the amino acids transferred by each tRNA, the corresponding anticodons, and the Ψ position on the tRNA molecules. **f** Venn plot shows the overlap of detected Ψ sites between biological replicates for *P. aeruginosa* and *P. syringae* during each growth phase.

To study Ψ modifications on bacterial RNA species beyond rRNA, we employed probe-based rRNA depletion(Choe et al., 2021) as a critical step in our optimized protocol for baBID-seq. We carefully established RNA fragmentation conditions to generate fragmented RNA of ∼60– 70 bp in length, ensuring efficient adaptor ligation; meanwhile, size selection of amplified library DNA by native PAGE gel effectively minimized contamination from adaptor dimers or dsDNA of unexpected sizes (**Supplementary Fig. 1a**). We then applied the baBID-seq protocol to four bacterial species (*K. pneumoniae, B. cereus, P. aeruginosa*, and *P. syringae*). Because the libraries were prepared from rRNA-depleted RNA, rRNA-derived signals may be variably affected across species. Accordingly, although residual 16S rRNA Ψ sites showed conserved modification fractions, we used these sites only as internal indicators of baBID-seq library performance in generating deletion signatures, rather than for comprehensive rRNA Ψ profiling, serving as benchmarks for assessing baBID-seq library quality (**Supplementary Fig. 1b**).

baBID-seq successfully captured various RNA species, including rRNA, tRNA, and mRNA. We characterized hundreds of Ψ sites on rRNA-depleted RNA isolated from four bacterial strains across exponential and stationary growth phases. The results from baBID-seq quantitatively demonstrated an excellent correlation between biological replicates (**Fig. 1c** and **Supplementary Fig. 2a**). While Ψ sites on rRNA and tRNA consistently showed stable modification levels across biological replicates, certain mRNA Ψ sites displayed greater variability.

Given the conservation of tRNA Ψ sites across biological replicates, we used the average Ψ fraction at each specific tRNA Ψ site for downstream analysis. baBID-seq quantitatively maps Ψ modifications at various positions within tRNA, including the stem and loop of the T-arm, the anticodon arm, and the D-arm (**Fig. 1d** and **Supplementary Fig. 2b,c**). To investigate tRNA Ψ dynamics during exponential versus stationary growth phases, we quantified Ψ fraction differences at each specific site across four strains. Our analysis revealed that most Ψ sites on bacterial tRNA consistently exhibited higher modification fractions in the stationary phase across all examined strains (**Fig. 1e**). In *K. pneumoniae*, the Ψ sites within the T-arm, D-arm, and anticodon arm collectively showed a reduced modification fraction in the exponential phase compared to the stationary phase (**Fig. 1d**). Similarly, in *B. cereus* and *P. syringae*, most tRNA Ψ sites within the T-arm displayed lower Ψ fractions during the exponential phase (**Supplementary Fig. 2b,c**). Previous research has shown that the T-arm of tRNA can globally influence tRNA maturation and regulate translation in *E. coli*(Schultz et al., 2024). Thus, this growth phase-dependent pattern of tRNA pseudouridylation suggests a coordinated regulatory mechanism that may affect mRNA translation as bacteria adapt to changing environmental conditions.

In addition to rRNA and tRNA, baBID-seq also identified highly conserved Ψ sites on mRNA between biological replicates (**Fig. 1f** and **Supplementary Fig. 2d**), providing strong confidence in the identification of genuine Ψ modifications. To further verify the site reliability, four Ψ sites were tested with pseU-TRACE(Fang et al., 2024): Ψ site at position 944 on 23S rRNA, a negative control site located within *guaA* gene, a Ψ site within *clpV1* gene, and an intergenic Ψ site located between *guaA* and *guaB* genes in *P. aeruginosa*. All three positive sites were successfully detected by pseU-TRACE, and no signal was observed at the negative-control site (**Supplementary Fig. 2e)**. For subsequent analysis, we focused exclusively on Ψ sites that were consistently detected across biological replicates.

### BID-seq uncovers abundant Ψ modification in bacterial mRNA

With the identification of highly conserved Ψ modifications in bacterial mRNA enabled by baBID-seq, we proceeded to analyze their distribution patterns and quantitative features across the transcriptome. In total, we detected over 3,000 Ψ sites in the mRNA of four bacterial strains. Notably, the metagene plot revealed that most Ψ sites were enriched within the coding sequences (CDS) of bacterial mRNAs, and this distribution pattern shows remarkable consistency across the four strains (**Fig. 2a,b**), which was also similar to the observations in human cell lines(Dai et al., 2023). Overall, the average Ψ modification in bacterial mRNA (mean Ψ fraction: 15%) was lower than that in 16S and 23S rRNA (mean Ψ fraction: 40%). Most mRNA Ψ sites were primarily distributed below a 25% Ψ fraction, while a smaller proportion reached modification fraction above 50% as highly modified ones (**Fig. 2c** and **Supplementary Fig. 3a**).

**Figure 2.**
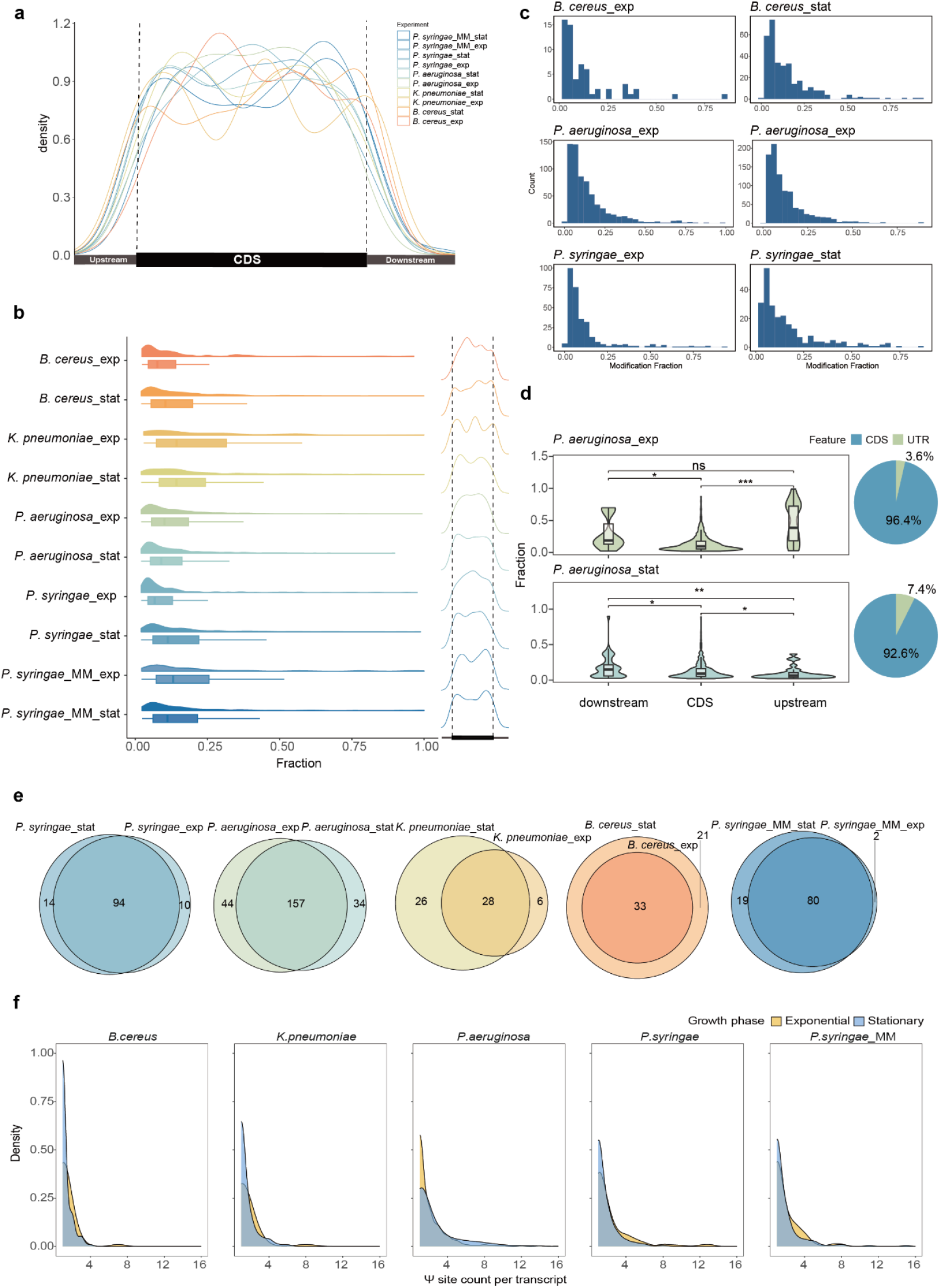
baBID-seq uncovers Ψ modification in bacterial mRNA CDS and UTRs. **a** Density plot depicting the distribution of Ψ modifications in mRNA across different growth phases and conditions. **b** Distribution of mRNA Ψ fraction showing strain and growth phase-specific patterns of Ψ distribution (right Ψ density plot across each strain’s mRNA). **c** mRNA Ψ faction and counts under different conditions. **d** Right pie charts show the proportion of Ψ sites in intergenic versus coding regions, and violine plots compare Ψ fraction values between untranslated regions (upstream and downstream UTRs) and coding regions. Statistical significance was determined using the Wilcoxon Signed-Rank test; ns, *p-*value ≥ 0.05; \**p-* value < 0.05; \*\**p-*value < 0.01; \*\*\**p-*value< 0.001 and \*\*\*\**p-*value< 0.0001. **e** The pie charts show the Ψ-modified gene overlap in two growth phases across four strains. **f** Density plot shows the Ψ site numbers per transcript.

Since Ψ modifications in mRNA untranslated regions (UTRs) can influence mRNA processing in eukaryotic cells(Martinez et al., 2022; Rodell et al., 2024), and the UTRs play a crucial role in post-translational regulation in bacteria(Adams et al., 2021), we conducted a detailed investigation of Ψ modifications in the upstream and downstream UTRs. We found that UTRs’ pseudouridylation accounted for 7% to 16.4% of total Ψ sites across the transcriptome in both stationary and exponential phases (**Supplementary Fig. 3b,c**). Notably, in *P. aeruginosa* and *B. cereus*, we observed a significantly higher modification fraction for Ψ sites within downstream regions compared to coding sequences (CDS) in both growth phases (**Fig. 2d** and **Supplementary Fig. 3d**). In contrast, *P. aeruginosa* exhibited a higher Ψ modification level in the upstream regions of mRNA during the exponential phase (**Fig. 2d**).

We defined a Ψ-modified gene as any mRNA containing one or more Ψ sites in any growth phase and identified both phase-shared and phase-unique genes across the four strains (**Fig. 2e**). We also noted that individual genes frequently harbored multiple Ψ sites, with the number of sites varying dynamically across bacterial growth phases (**Fig. 2f**).

### Motif analysis of Ψ modifications in bacterial mRNA, rRNA and tRNA

Previous research has shown that different RNA species exhibit unique Ψ modification patterns, with Ψ synthases targeting specific sequence motifs in tRNA and rRNA (such as the RluA motif ΨURAA)(Pan et al., 2003; Schaening-Burgos et al., 2024). To identify the sequence determinants of Ψ modifications and uncover RNA-type-specific Ψ motif contexts across bacterial transcriptomes, we conducted a comprehensive motif analysis of Ψ-modified sites with a fraction above 2%, which could be reliably identified in both growth phases. The sequence context analysis focused on 5-nucleotide motifs centered on each Ψ site. We calculated the percentages of Ψ motifs by dividing the count of each unique Ψ motif by the total number of Ψ motifs detected in the mRNA of a single bacterial strain (**Fig. 3a**).

**Figure 3.**
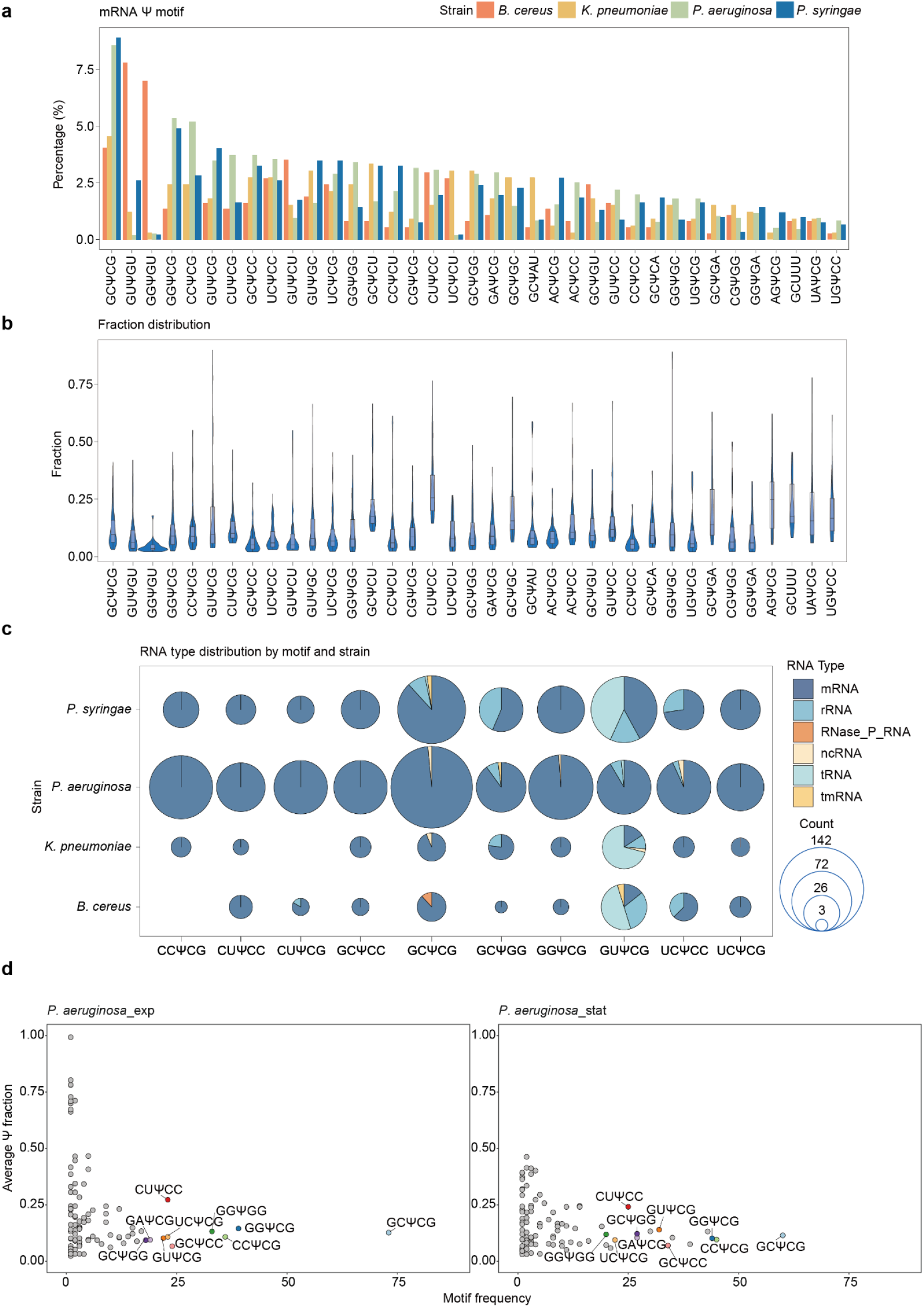
Comparative analysis of Ψ modification motif across strains. **a** Comparison of overall mRNA 5-mer Ψ motif ratios across four strains. Motif ratios are calculated by dividing the count of each specific 5-mer motif centered on Ψ by the total number of motifs detected in each individual strain mRNA. Ψ-modified sites with a fraction above 2% are used here. **b** Distribution of Ψ fraction (ranging from 2% to 100%) for each motif detected. **c** Scatter pie chart shows the proportional distribution of top 10 (ranked by motif abundance) Ψ-containing motif counts categorized by RNA types and bacterial strains. **d** The scatter plot illustrates the relationship between the average modification fraction and abundance of motifs in *P. aeruginosa* in exponential (*P. aeruginosa*_exp) and stationary (*P. aeruginosa*_stat) growth phases. The average Ψ fraction was calculated as the sum of Ψ fractions for each individual motif divided by its frequency.

Our analysis revealed a diverse array of Ψ motifs within bacterial mRNA based on baBID-seq data. Notably, GCΨCG, GGΨCG, and CCΨCG were the most abundant motifs observed in the three Gram-negative bacteria, while GUΨGU and GGΨGU were the dominant motifs in *B. cereus* (**Fig. 3a**). We then analyzed the quantitative features of Ψ sites within these diverse motif contexts and found that the average Ψ fractions for different motifs ranged from 3.4% to 96.6%, indicating varying Ψ installation efficiencies regulated by the properties of different Ψ synthases (**Fig. 3b**). Overall, we summarized the top 10 frequent Ψ motifs for bacterial mRNA of *P. aeruginosa*, and *P. syringae*: GUΨCG, (CC/CU/GC/GG/UC)ΨCG, (CU/GC/UC)ΨCC and GCΨGG (**Fig. 3c**).

baBID-seq also reveals Ψ motif contexts in bacterial rRNA and tRNA. While a variety of Ψ motifs were identified in bacterial rRNA, the GUΨCG motif stands out as the predominant Ψ motif in tRNA across the four bacterial strains (**Supplementary Fig. 4a–d**). Notably, the GUΨCG motif is well-characterized within the T-arm of tRNAs (at position 55) and is specifically modified by TruB family(Dai et al., 2023; De Crécy-Lagard et al., 2019; Hoang and Ferré-D’Amaré, 2001; Pan et al., 2003; Schultz et al., 2024; Veerareddygari et al., 2016). According to our baBID-seq data, GUΨCG has been confirmed as the predominant motif in both tRNA and rRNA (mean fraction: 68%), as well as in mRNA (mean fraction: 19%) (**Fig. 3c** and **Supplementary Fig. 4c,d**), suggesting that TruB may also play a role in Ψ installation in bacterial mRNA. We also identified several other key motifs, including UUGC, UUGA, and UUAAA, which correspond to the previously characterized RluA motif in *E. coli*(Schaening-Burgos et al., 2024). Overall, no distinct sequence motifs were universally enriched in the mRNA of the four bacterial strains (**Supplementary Fig. 4e–h**), likely due to the complex interactions among multiple Ψ synthases involved in U-to-Ψ conversion on mRNA.

To determine whether sequence preferences for Ψ modifications vary under different growth conditions, we calculated both motif frequency and average Ψ fraction for each Ψ motif in *P. aeruginosa*. Notably, we observed a decrease in the frequency of the GCΨCG motif (**Fig. 3d**), as the most abundant Ψ motif among the three Gram-negative bacteria. For other top Ψ motifs, the associated modification levels showed only slight variations between the exponential and stationary phases. To gain insights into the sequence features related to Ψ modifications, we analyzed the nucleotide composition within 10 base pairs flanking Ψ sites in bacterial mRNA. While most strains exhibited non-unique differences in GC content, *B. cereus* displayed a notably higher GC ratio (normalized to the genomic background GC content) compared to the other three bacterial strains (**Supplementary Fig. 4i–k**). To explore the relatively distinct motif pattern in *B. cereus*, we conducted comparative orthology-based analyses to identify pseudouridine synthase sequences among the five bacterial strains (**Supplementary Fig. 4l**). A unique interactive regulation of pseudouridine synthases in *B. cereus* may explain its divergent Ψ sequence patterns compared to other bacterial strains.

### Evolutionary conservation of clustered Ψ modifications in bacterial orthologous genes

To investigate the evolutionary conservation of mRNA Ψ modifications among bacterial strains, we analyzed orthologous genes, focusing on both the preservation of modification sites and the functional characteristics of Ψ-modified genes. Based on clustered Ortho groups and genome annotations, our results revealed 225 homologous genes carrying Ψ modifications in at least two bacterial strains (**Fig. 4a**). We systematically characterized the biological functions of these homologous Ψ-modified genes among the four bacterial strains through pathway enrichment analysis, using annotated gene sets from the Kyoto Encyclopedia of Genes and Genomes (KEGG). This analysis uncovered three distinct but interconnected metabolic clusters. The primary cluster showed highly significant enrichment in central carbon metabolism, including glycolysis, TCA cycle, oxidative phosphorylation, and amino acid biosynthesis (**Fig. 4a** and **Supplementary Fig. 5a**). Notably, gene clusters essential for ATP production, such as *atpA* and *atpD*, were also enriched. Among the four bacterial strains, the functional enrichment of Ψ-modified genes in core metabolic pathways, combined with the conservation of Ψ motif contexts (**Fig. 3a**), provides evidence for the functional importance and evolutionary conservation of a specific group of Ψ-modified transcripts across bacterial systems. In addition to the homologous Ψ-modified genes, we identified 633 strain-specific Ψ-modified mRNAs, highlighting the dynamic nature of Ψ modifications and suggesting potential strain-specific regulatory mechanisms.

**Figure 4.**
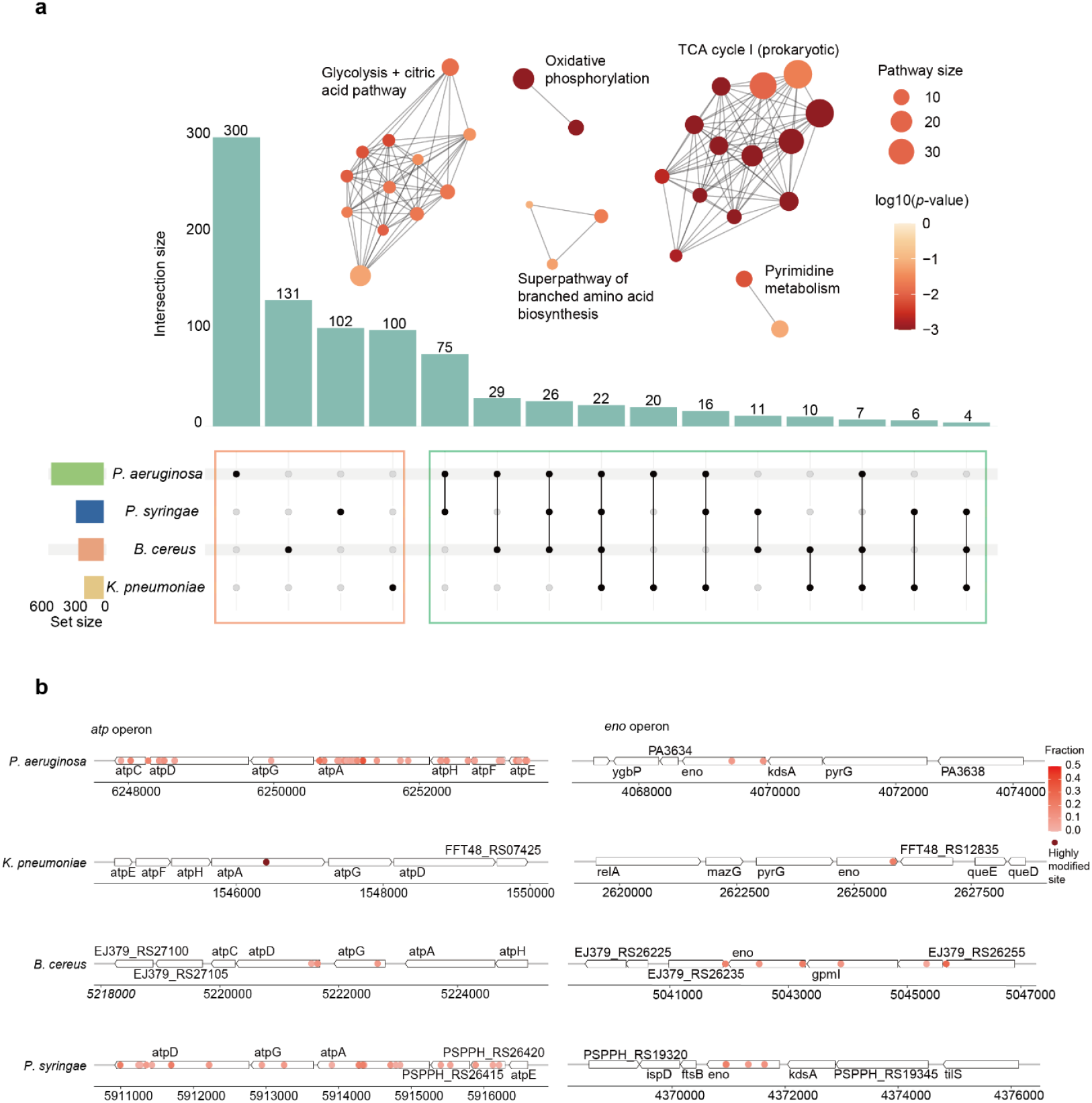
Evolutionary conservation of clustered Ψ modifications in orthologous genes. **a** The bar plot depicts the Ψ modified homologous genes among bacterial species (green box) and strain-specific genes (orange box). Functional networks generated by clustering KEGG pathway enrichment results of Ψ-modified homologous genes present across two or more strains. The dot size in the network indicates the gene number contained in specific KEGG pathways. The *p*-value for each pathway was calculated with Fisher’s Exact test. **b** Regions with Ψ enrichment across the *atp* and *eno* operons in four strains. Dark red dots represent highly modified sites with Ψ fraction values exceeding 50%.

We then conducted a functional investigation of Ψ-modified genes in the two growth phases of *P. aeruginosa*. The Gene Ontology (GO) enrichment analysis revealed that Ψ-modified genes were significantly enriched in GO terms related to energy metabolism during the exponential phase compared to the stationary phase, highlighting distinct state-specific GO patterns (**Supplementary Fig. 5b**). The exponential phase is known to exhibit elevated levels of metabolism-related genes, reinforcing our findings regarding the biological relevance of Ψ modifications. Notably, Ψ-modified genes showed a preferential enrichment in type IV swarming motility during the stationary phase. For instance, the key transcription factor *algR*, which regulates multiple virulence factors and promotes *P. aeruginosa* twitching motility, serves as a representative example among the Ψ-modified genes(Kong et al., 2015). This finding suggests a specific role for Ψ modifications in coordinating motility behaviors during the stationary growth phase, potentially linking RNA modification to adaptive virulence mechanisms when nutrient availability is limited. Besides, we observed that homologous genes exhibited more stable Ψ fraction patterns, likely due to their enriched functions in fundamental metabolic processes (**Supplementary Fig. 5c**).

To explore the regional distribution of Ψ modifications in greater detail, we conducted a 500 bp sliding window analysis across all detected mRNA transcripts in four bacterial strains, identifying specific regions with multiple Ψ sites using baBID-seq (**Supplementary Fig. 5d**). Further analysis revealed that most Ψ-enriched regions corresponded to operons or gene clusters. We identified Ψ clusters containing two or more Ψ sites in operons such as the evolutionarily conserved *atp* operon(Ventura et al., 2004) and *eno* operon, across all strains(Commichau et al., 2009; Salgado et al., 2000) (**Fig. 4b**). This phenomenon was also observed in other homologous gene clusters or operons, such as *rpoA, fusA, groE*, and *rpc* operons(Salgado et al., 2006) (**Supplementary Fig. 5e**), suggesting that Ψ modification may play a role in regulating post-transcriptional processes(Rodell et al., 2024). Upon examining the Ψ distribution within operons and gene clusters in more detail, we observed distinct regional patterns of Ψ modifications. Some Ψ modifications accumulated within the 3’ UTR regions of *atp* operon, for instance, in *atpD* of *P. aeruginosa* (**Fig. 4b**). In *P. aeruginosa*, Ψ sites in the *atpD* gene were predominantly located near stop codon, whereas in *B. cereus*, most Ψ modifications were concentrated at the translation initiation region (**Fig. 4b**). This detailed regional analysis of Ψ modifications within gene clusters revealed strain-specific distribution patterns in the conserved operons. The flexible regional Ψ modification patterns observed in operons such as *atp* may regulate gene expression to align with bacteria-specific metabolic needs.

### Growth state-dependent dynamic Ψ modification in bacterial mRNA

Ψ modification levels on mRNA have been reported to fluctuate under stress conditions in human cells(Li et al., 2015). Our study also observed alterations in Ψ modifications across the four bacterial strains under different growth conditions. We conducted a detailed comparative analysis of Ψ sites between the two growth phases, identifying both newly emerged and diminished Ψ modification events, as well as alteration in modification fractions at conserved sites. The quantitative baBID-seq approach allowed us to pinpoint dynamic Ψ modifications in response to bacterial metabolic shifts and changes in growth states. We initially compared the distribution of modification fractions for all mRNA Ψ sites in the exponential phase versus the stationary phase. *K. pneumoniae* and *B. cereus* exhibited significantly higher global Ψ levels during the stationary phase (**Supplementary Fig. 6a,b**). In contrast, no matter in nutrient-enriched or minimal media (MM) conditions, *P. aeruginosa* and *P. syringae* did not show significant changes in global Ψ fractions between the two phases (**Supplementary Fig. 6c,d**).

We then focused on changes in Ψ fractions at specific sites, setting a cutoff of 10% variation between the two growth phases across the four bacterial strains. In *P. aeruginosa* and *P. syringae*, we identified numerous phase-specific Ψ sites that either emerged or disappeared, as well as many conserved Ψ sites exhibiting changes in Ψ fractions above 10% (**Fig. 5a,b**). For instance, in *P. aeruginosa*, key genes linked to energy metabolism, such as *sucB, sucC*, and *gltA*, displayed Ψ sites of increased modification fraction during the exponential phase of heightened metabolic activity. The stop codon region of *secY*, a protein essential for type II secretion systems in *P. aeruginosa*, contained a highly modified Ψ site that was only detected during the exponential growth phase; similar Ψ fraction dynamics were observed for *secY* in *K. pneumoniae* and *B. cereus* (**Supplementary Fig. 6a–c**).

**Figure 5.**
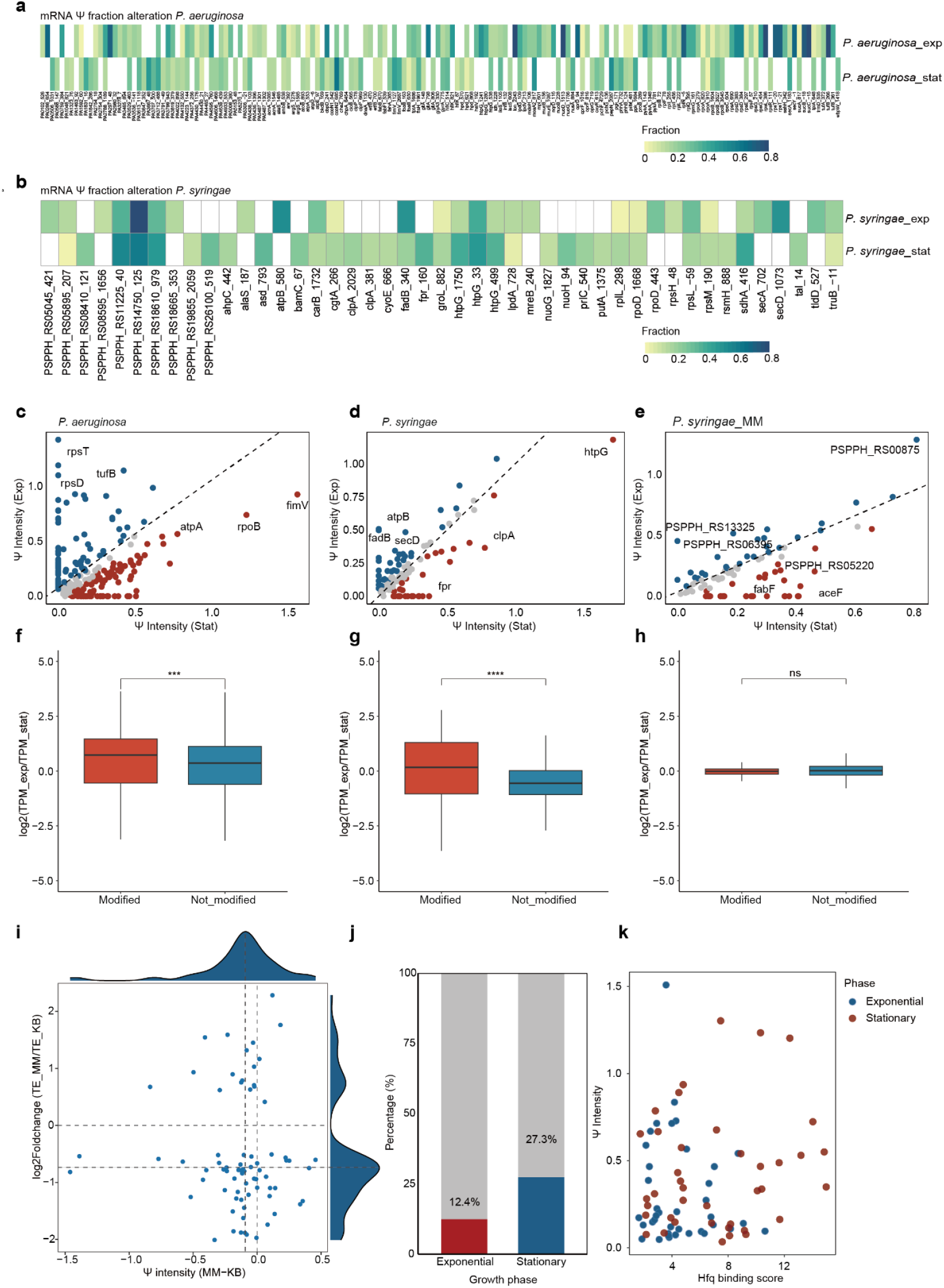
Growth state-dependent dynamics of Ψ modification fraction. **a**,**b** Heatmap showing the mRNA Ψ fractions alteration of each site in *P. aeruginosa* (a) and *P. syringae* (b). The color intensity reflects Ψ fraction at each site. Only sites with >10% absolute difference in Ψ fraction between exponential and stationary phases are displayed. Blank boxes signify either unmodified sites or those with Ψ fractions below 2%. The annotation combining position label and gene name indicates the precise location of Ψ modification within genes. **c** Scatter plot shows Ψ intensity alteration between two growth phases in *P. aeruginosa*. For each mRNA, Ψ intensity is calculated as the sum of all Ψ fractions throughout the transcript. **d** Similar to **c**, the scatter plot shows Ψ intensity alteration in *P. syringae* under two growth phases. **e** scatter plot shows Ψ intensity alteration in *P. syringae* under two growth phases in MM medium condition. **f-h** Box plot shows the modified and unmodified mRNA transcripts per kilobase million (TPM) changing between exponential and stationary growth phases under different conditions: *P. aeruginosa* cultured in LB medium (**f**), *P. syringae* cultured in KB medium (**g**) and *P. syringae* cultured in MM medium (**h**). Y-axis shows log2(TPM at exponential phase / TPM at stationary phase) of each mRNA. The red color presents Ψ-mRNA, and the blue color indicates no-Ψ-mRNA. Wilcoxon Signed-Rank test; ns, *p-*value ≥ 0.05; \**p-*value < 0.05; \*\**p-*value < 0.01; \*\*\**p-*value< 0.001 and \*\*\*\**p-*value< 0.0001. **i** The scatter plot illustrates the correlation between translation efficiency (TE) alteration (log2Foldchange of (TE_MM/TE_KB)) and Ψ intensity difference (Ψ intensity of MM-Ψ intensity of KB) in *P. syringae* cultured under MM medium (TE_MM) versus KB conditions (TE_KB). **j** The proportion of Hfq-bound Ψ-mRNA (red color for exponential growth phase mRNA and blue color for stationary phase mRNA) versus those non-Ψ-mRNA (grey color) across two growth conditions in *P. aeruginosa*. **k** The scatter plot shows a correlation between mRNA Ψ intensity and Hfq binding score, where the Hfq binding score is calculated as the sum of each mRNA peak binding strength (log2FoldChange value of each peak).

In addition to analyzing Ψ sites, we also calculated Ψ intensity for each gene, defined as the sum of modification fractions for all Ψ sites within a single mRNA molecule. During the transition from the exponential to the stationary phase, the Ψ intensity of specific genes shifted significantly (**Fig. 5c–e** and **Supplementary Fig. 6e**). In the exponential phase, many genes involved in metabolism, amino acid biosynthesis, and protein synthesis exhibited higher Ψ intensity, such as *rpsD, rpsT, tufB, lon, nuoD*, etc., reflecting their critical role in supporting rapid growth; meanwhile, for these genes, their decreased Ψ intensity observed in the stationary phase may suggest a coordinated Ψ reprogramming that helps bacteria adapt to reduced nutrient availability and increased cell density by downregulating energy-intensive processes. Overall, the dynamic nature of Ψ modifications—including newly emerged and disappeared Ψ sites, as well as altered Ψ fractions across the two growth conditions—indicates their potential roles as responsive epi-transcriptomic switches that facilitate bacteria to be adapted to varying environmental challenges and metabolic demands.

### Ψ correlates with bacterial mRNA metabolism and function

It is well established that Ψ can enhance mRNA stability and translation in mammals(Karikó et al., 2008), parasites(Li et al., 2025; Nakamoto et al., 2017), and plants(Li et al., 2025). However, the effects of Ψ on bacterial mRNA have yet to be investigated. We normalized gene expression levels by calculating transcripts per kilobase million (TPM) for the two growth phases and grouped the adequately expressed genes into Ψ-modified mRNA (Ψ-mRNA) and non-Ψ-modified mRNA (non-Ψ-mRNA). The analysis revealed that the expression level of Ψ-mRNA was significantly higher than that of non-Ψ-mRNA in *P. aeruginosa* and *P. syringae* when cultured in a nutrient-sufficient medium (**Fig. 5f,g** and **Supplementary Fig. 6f**). However, *P. syringae* exhibited more moderate changes in mRNA expression between the different growth phases under minimal medium (MM) conditions (**Fig. 5h**). Our findings suggest that Ψ may stabilize mRNA in a growth phase-dependent manner in bacteria.c

We then conducted comprehensive analyses of TE in *P. syringae* with alterations in Ψ modifications under two distinct conditions (King’s B medium, KB and minimal medium, MM) to examine whether Ψ impacts bacterial mRNA translation. Although we did not observe a universal global correlation between changes in Ψ modifications and TE for all genes under both conditions, a larger proportion of genes did show a positive correlation between TE and Ψ modifications (**Fig. 5i**). Our findings partially align with previous studies in mammals, which suggested that Ψ modifications could enhance mRNA translation efficiency in bacteria and may play more complex roles in translation regulation.

Hfq is a major bacterial post-transcriptional regulator that functions as a pivotal RNA-binding protein (RBP), orchestrating various cellular processes(Trouillon et al., 2022). Its regulatory mechanisms have been extensively characterized, including the alteration of RNA structure and the facilitation of sRNA-mRNA interactions(Chihara et al., 2019), highlighting its fundamental role in coordinating gene expression networks (dos Santos et al., 2019; Sobrero and Valverde, 2012). To investigate the potential association between Hfq and Ψ modifications, we performed an integrative analysis combining our data with previously published Hfq RIP-seq data(Trouillon et al., 2022) in *P. aeruginosa*. We examined both exponential and stationary growth phases to evaluate whether Hfq targets Ψ-modified regions or whether Ψ modifications influence Hfq-RNA interactions. Our results indicated that the distance between Ψ sites and Hfq peak centers significantly decreased during the stationary phase (**Supplementary Fig. 6g**); meanwhile, Hfq-bound genes accounted for a substantial proportion of mRNA Ψ sites, with 12.5% during the exponential phase and 27.3% during the stationary phase (**Fig. 5j**). By defining the Hfq binding score as the sum of Hfq peaks binding strength per gene, we found that, in the stationary phase, Hfq exhibited a stronger binding affinity for genes carrying more Ψ modifications (**Fig. 5k**). These results suggest that Ψ modifications may facilitate mRNA-Hfq interactions to some extent. Overall, dynamic Ψ modifications have been shown to impact bacterial mRNA stability, translation, and RBP interactions in response to changing cellular demands during growth phase transitions.

### Integrated computational analysis reveals structure-dependent Ψ modifications in bacterial RNA

Although some Ψ modifications shared common sequence motifs, we observed an array of diverse motif contexts at Ψ sites in bacterial mRNA (**Fig. 3a,b**). This observation aligns with previous studies demonstrating that pseudouridine synthases, such as PUS1 and TruB, preferentially recognize RNA local structures beyond primary sequence motifs for Ψ installation(Carlile et al., 2019; Lange et al., 2012; Pan et al., 2003; Safra et al., 2017). To model and study the widespread Ψ modifications on bacterial mRNA, which are hypothesized to be structure-dependent, we incorporate Ψ modification determinants through comprehensive clustering analyses of local RNA sequences and structural elements to test our hypothesis. We firstly calculated the predicted secondary structure 41nt centered GUΨC motif, with the representative Ψ sites of 50%∼96% modification fraction (**Fig. 6a**). Interestingly, all GUΨC motifs with varying Ψ fractions are predicted to occur within RNA loop structures. To determine whether RNA structural factors influence Ψ deposition and modification fractions, we compared the structural and fractional characteristics of the two highly prevalent motifs, GUΨC and GCΨCG, by clustering predicted RNA structures across all RNA species in *P. aeruginosa*. The results indicated that, aside from tRNA and rRNA, which clustered together due to their distinct structural features, we observed small clusters within certain mRNAs, such as *guaB, recA*, and *PA4943*. Overall, we did not find characteristic structural signatures that could completely distinguish GUΨC from the GCΨCG motif (**Fig. 6b**). To gain deeper insights, we conducted structural clustering analyses of all RNA species across different strains (**Supplementary Fig. 7a-d**). Given that some pseudouridine synthases target specific structural features, we anticipated highly distinguishable clustering results. However, contrary to our expectations, such distinct clustering patterns were not observed. This may suggest that certain pseudouridine synthases, such as RluA(Schaening-Burgos et al., 2024), do not solely rely on structural features for target recognition. Notably, we identified clusters of Ψ sites with similar structures that exhibited higher modification fractions, including *sucC* and *sucB* in *P. aeruginosa* (**Supplementary Fig. 7a**), as well as *sdhA, PSPPH_RS14750*, and *atpB* in *P. syringae* (**Supplementary Fig. 7c**). Integrating these findings with the distinctive Ψ fraction patterns exhibited by various motifs (**Fig. 3b**), we conclude that both local sequence and structural features of RNA may play important roles in Ψ installation.

**Figure 6.**
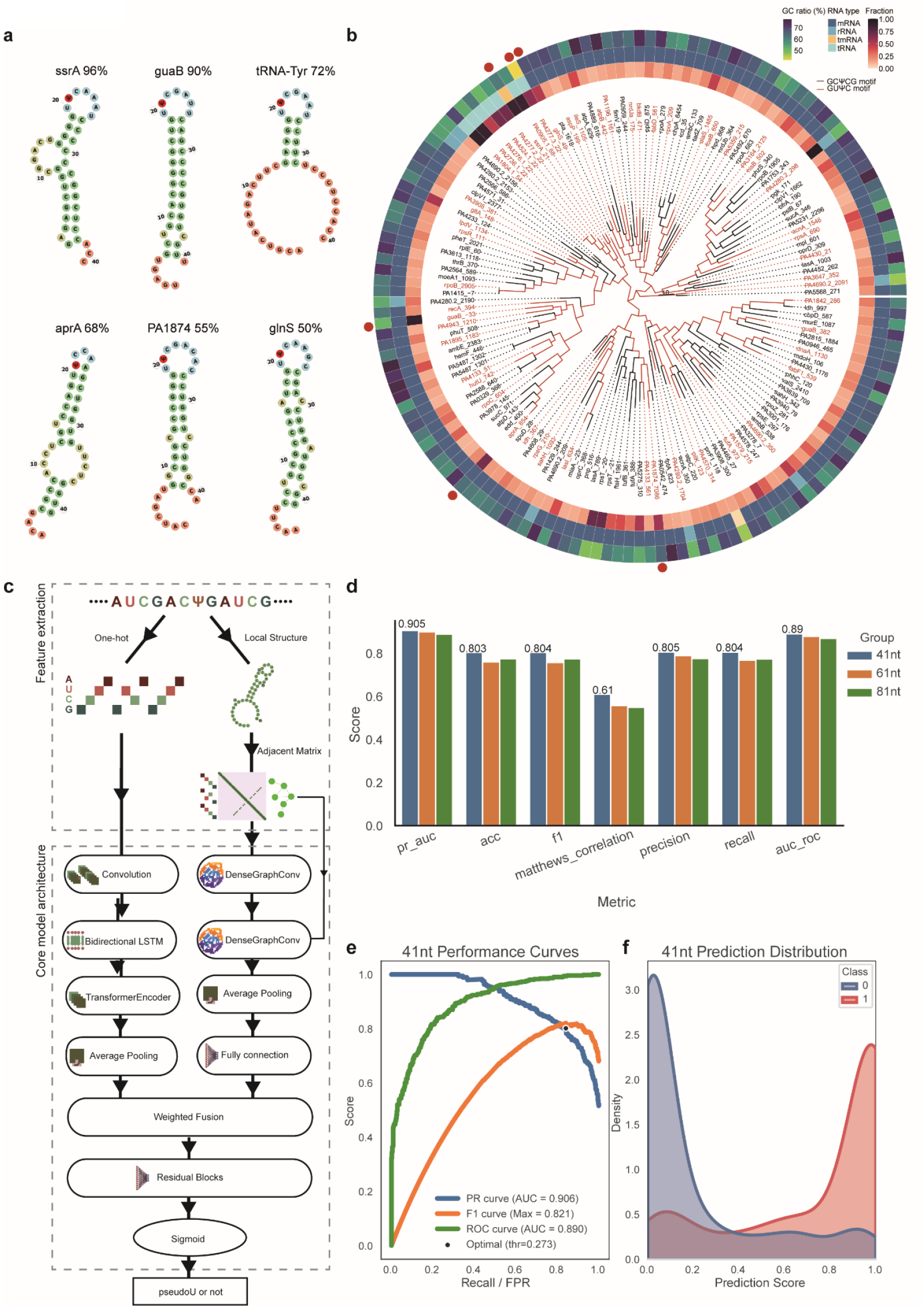
Structure-dependent patterns of Ψ modifications and transformer-GNN-based deep learning network for Ψ prediction. **a** Predicted RNA secondary structures containing the GUΨCG motif with corresponding Ψ fraction values and gene identifiers annotated. MXfold2 is employed to model these structures using 20nt flanking sequences extending from each modification site. **b** Sequence and structure clustering of 41-nucleotide RNA segments centered by Ψ sites with fraction values greater than 0.1 and containing either GCΨCG (black branch color) or GUΨC (red branch color) motifs. The circular visualization features three concentric layers: the inner layer displays the Ψ fraction value, the middle layer indicates RNA type, and the outer layer represents the GC ratio (%) of each 41nt RNA segment. The red dot around the circle marked the position of RNA displayed in (**a**). **c** Architecture of pseU_NN. The model integrates sequence and structural information through two parallel pathways: 1) A sequence analysis branch with one-hot embedding followed by a multi-head transformer module, and 2) A structure analysis branch that processes RNA secondary structure adjacent matrices through a graph convolution module to extract structural features. The features extracted by the two modules are further weighted and merged as input for residual blocks (fully connected layers). **d** Bar plots summarize model performance with input sequences of 41 nt, 61 nt, and 81 nt, evaluated by PR-AUC, accuracy (ACC), F1 score, Matthews correlation coefficient (MCC), precision, recall, and ROC-AUC. Overall, the three sequence lengths show comparable performance across metrics, with the 41 nt model achieving slightly higher PR-AUC and ROC-AUC, indicating that shorter sequence contexts are sufficient for robust pseudouridine prediction. **e** Multi-metric assessment showing precision-recall curve (AUC 0.906), F1 curve (AUC 0.821), and ROC curve (AUC 0.89) of the pseU_NN model on 41nt validation datasets, achieving a peak F1 score of 0.804. **f** Distribution of pseU_NN prediction scores on 41nt test datasets.

### LSTM-Transformer-GNN-based neural networks for integrated prediction of Ψ

In next-generation sequencing (NGS) data, variable read coverage dictated by gene expression patterns or limited sequencing depth can lead to missed potential Ψ sites. To address this limitation, we implemented a strategy that integrates RNA sequence and structural features for a transcriptome-wide scan of Ψ sites, resulting in a more comprehensive inventory of potential Ψ sites. Previous studies have shown that the sequence context surrounding Ψ sites can serve as a reliable predictor for RNA modifications(Hoang and Ferré-D’Amaré, 2001; Song et al., 2021). Building on this, we developed a deep learning model that accurately captures both sequence and structural features of known Ψ sites across various RNA species, allowing us to predict potential modification sites that may be condition-dependent or below the detection thresholds of BID-seq. We extracted sequence segments of 41, 61, and 81 nucleotides (±20, ±30, and ±40 bp) centered on each Ψ site, applying window shifts of ±5, ±10, and ±15 bp, respectively. The input sequences were then embedded using one-hot encoding and processed through a multi-head transformer module after the convolution layers and the bidirectional LSTM (Long Short-Term Memory) layer. Simultaneously, we utilized adjacency matrices representing local RNA structures (predicted using MXfold2(Sato et al., 2021) as input for a graph neural network (GNN) module. Features extracted from both modules were combined through weighted concatenation and subsequently processed using a residual block. We employed binary cross-entropy loss for predicting the likelihood of Ψ modifications. This hybrid LSTM-transformer-GNN architecture effectively integrated both sequence and structural characteristics across various RNA sequences (**Fig. 6c**), termed as pseU_NN.

We used 3,377 high-confidence Ψ sites (with fraction values >2%) as positive samples. The negative samples consisted of 3,400 randomly selected sites that contained the unique Ψ motif but showed no evidence of Ψ deposition in our experimental data. The dataset was divided into 4,744 training samples, 1,016 test samples, and 1,017 validation samples. The model consistently performed well across different input dimensions (41, 61, and 81 nucleotides), with all variants achieving AU-ROC scores exceeding 0.8 after convergence (**Fig. 6d**). Using 41-nucleotide inputs, our methodology achieved impressive validation metrics, including an area under the precision-recall curve (AU-PRC) of 0.905 and an area under the receiver operating characteristic curve (AU-ROC) of 0.89 (**Fig. 6e**). Models trained with alternative input sequence lengths also demonstrated strong performance metrics (**Supplementary Fig. 7e-h**). These results set the basis for further development of effective deep learning tools for transcriptome-wide Ψ prediction in both bacterial and mammalian RNAs.

## Discussion

RNA modifications in bacteria, particularly Ψ, are less characterized than their well-studied eukaryotic counterparts. Leveraging recent advances in quantitative sequencing methods such as BID-seq(Dai et al., 2023; Zhang et al., 2024), we present comprehensive, single-nucleotide resolution maps of Ψ modifications, complete with stoichiometric information, across four bacterial strains under different growth phases. Our findings confirm the widespread occurrence of Ψ modifications in bacterial RNA and provide a functional analysis of these modifications. This extensive dataset serves as a valuable resource for understanding the evolutionary and functional significance of RNA Ψ modifications in bacteria.

The Ψ modification in tRNA plays critical regulatory roles in tRNA aminoacylation, stability, and the formation of functional structures(De Crécy-Lagard et al., 2019; de Crécy-Lagard and Jaroch, 2021; Krutyhołowa et al., 2019; Schultz et al., 2024). Our analysis revealed that tRNA Ψ modification is present in varying fractions, with a stronger modification level observed in the TΨC loop compared to the anticodon-arm and D-arm loops. Previous studies indicate that the tRNA T-arm is highly modified, not only by Ψ but also by other uridine modifications like 5-methyluridine (m^5^U)(Chou et al., 2017). Both Ψ and m^5^U modifications globally enhance tRNA aminoacylation, and also independently influence specific tRNA modifications, such as 3-(3-amino-3-carboxypropyl)uridine at position 47(Schultz et al., 2024). The TΨC loop is crucial for the interaction between tRNA and the ribosome, facilitating the formation of the tRNA-ribosome complex(Chou et al., 2017). Interestingly, we observed a novel phenomenon where Ψ modifications on the tRNA T-arm increase in the stationary growth phase compared to the exponential phase. Given that codon composition and mRNA expression are closely correlated(Gouy and Gautier, 1982), the dynamic Ψ modification in the TΨC loop may regulate bacterial mRNA translation by modulating T-arm interactions with the ribosome. Besides, other RNA modifications on tRNA may be influenced by the dynamic Ψ pattern during growth phase transitions, potentially creating a feedback loop where existing modifications affect the biogenesis of subsequent modifications. Overall, this condition-dependent Ψ modification in tRNA may represent a new mechanism by which bacteria adapt to their environment, anticipating further investigation in future research.

Previous studies have demonstrated that both a deficiency and an excess of pseudouridine can severely impair ribosomal translation and proper assembly in *E. coli*(Leppik et al., 2017; O’Connor et al., 2018). The stable fraction of Ψ modification observed in *E. coli* rRNA across two different growth phases suggests that rRNA Ψ may be tightly regulated to maintain essential rRNA function. The role of Ψ in mRNA remains unclear across the three domains of life. Our results reveal a comprehensive mRNA Ψ landscape in four bacterial strains. We found that the overall fraction of mRNA Ψ modifications was significantly lower than that of rRNA and tRNA, consistent with findings in plants and mammals(Dai et al., 2023; Li et al., 2025). The distribution and stoichiometric patterns of mRNA Ψ between Gram-positive and Gram-negative bacteria exhibited similarities. Notably, we identified evolutionarily conserved Ψ modifications in mRNAs encoding proteins involved in energy generation, ATP binding, amino acid synthesis, and protein translation, mirroring observations in mammals and plants(Dai et al., 2023; Li et al., 2025). We also discovered clusters of multiple Ψ sites enriched in specific operons related to conserved functions, which were detected across multiple strains.

To date, limited research has focused on Ψ modifications and their alterations in bacteria. In this study, we profiled and uncovered dynamic changes in Ψ modifications during growth phase transitions among four bacterial strains. During the metabolically active phase of *P. aeruginosa*, we observed increased Ψ modifications in many metabolism-related genes (**Supplementary Fig. 5b**). In the stationary phase, our analysis revealed reduced pseudouridylation within the coding regions of *fimV*, a gene that encodes an inner membrane protein in *P. aeruginosa* responsible for regulating intracellular cyclic AMP levels, type IV-mediated twitching motility, and type II secretion system genes(Buensuceso et al., 2016; Semmler et al., 2000). Ψ modifications in coding regions are known to alter codon properties on mRNA, leading to reconstituted translation and promoting the low-level synthesis of multiple peptides(Eyler et al., 2019). These findings suggest a potential new mechanism for regulating bacterial metabolism and quorum sensing in *P. aeruginosa*. Previous studies have demonstrated that pseudouridylation enhances mRNA stability in mammals and plants(Dai et al., 2023; Li et al., 2025; Zhang et al., 2024), and our research confirms and extends these observations to bacterial systems.

It has been reported that methionine aminoacyl tRNA^Met^ synthetase can target Ψ1074 in yeast (Levi and Arava, 2021), and several Ψ sites overlap with RNA-binding protein (RBP) binding regions(Martinez et al., 2022). To systematically investigate dynamic Ψ modifications in bacterial RNA and their potential impact on RBP binding, we conducted an integrative analysis that combined our BID-seq data with Hfq RIP-seq data from *P. aeruginosa*. Hfq is an RNA chaperone that recognizes 5-repeat AAN motifs in *P. aeruginosa* and plays a crucial role in regulating various post-transcriptional processes, including mediating sRNA-mRNA interactions(Chihara et al., 2019). Hfq can trigger mRNA structural reprogramming, and RNA structural switches may facilitate or hinder the process of pseudouridylation(Carlile et al., 2019; Hua et al., 2024). We observed increased Hfq binding to Ψ-modified mRNAs during the stationary phase compared to the exponential phase, suggesting that Ψ may modulate Hfq-mediated regulation via direct effects on RNA–protein affinity and/or indirect remodeling of mRNA structure; however, to establish causality, targeted perturbation of specific pseudouridine synthases and in vitro Hfq–RNA interaction assays with defined Ψ-containing versus unmodified transcripts are needed to confirm the role of Ψ in regulating Hfq–RNA interactions.

Ψ modification is specifically recognized in RNA partial structures or local motif contexts by the TruB pseudouridine synthase(Machnicka et al., 2014). Our comprehensive analysis of Ψ-containing sequences in mRNA revealed conserved motif signatures that closely resemble those found in tRNA and rRNA. This conservation pattern suggests that tRNA and rRNA Ψ synthases may directly recognize similar structural and sequence elements in mRNA, as evidenced by PUS1 and PUS6, which can both add Ψ to tRNA and mRNA(Carlile et al., 2019; Levi and Arava, 2021). Through an integrated approach combining RNA sequence and structural clustering analysis, we demonstrated that both the selection of Ψ sites and the modification fraction are strongly associated with the local secondary structure of RNA molecules(Carlile et al., 2019). From a structural perspective, this specificity likely explains why certain potential modification sites with appropriate motifs remain unmodified or exhibit dynamic Ψ levels under different growth conditions.

Deep learning methods are increasingly utilized for RNA modification prediction. Building on previous concepts, such as attention-based multi-label neural networks that predict multiple RNA modifications using sequence context(Song et al., 2021), we developed a hybrid LSTM-transformer-GNN architecture that integrates RNA structural features with multi-head attention mechanisms to predict potential Ψ modification sites (pseU_NN), particularly those that may have eluded detection due to low or absent gene expression. The robustness of our model was rigorously validated using high-confidence sites of various lengths and predicted RNA structures as input. pseU_NN enables reliable prediction of potential Ψ sites across different bacterial contexts, providing a comprehensive map of potential Ψ sites within bacterial transcriptomes.

Overall, this study presents the first comprehensive landscape of Ψ modifications across multiple diverse bacterial strains using optimized BID-seq. We systematically revealed Ψ fractions in tRNA, rRNA, and mRNA under exponential and stationary growth conditions in four strains, as well as under nutrient-deficient conditions in *P. syringae*. Our findings indicated that Ψ modifications on rRNA were stable under both growth conditions in *E. coli* and *P. aeruginosa*, while significant variations in Ψ fractions were observed in tRNA and mRNA across different growth phases. The motif conservation analysis provides insights into the activity of pseudouridine synthases on bacterial mRNA. We further identified evolutionarily conserved patterns of Ψ-enriched modifications in conserved operons involved in fundamental metabolic pathways across bacterial strains. In summary, our comprehensive study enhances the understanding of the regulatory roles of Ψ modifications across the bacterial kingdom, paving the way for future mechanistic investigations into the unknown functions of Ψ in bacterial RNA regulation.

## Materials and Methods

### Bacteria strains and growth conditions

The wild-type *Pseudomonas syringae pv. phaseolicola* 1448A strain was cultured in King’s B (KB) medium(King et al., 1954) (20 g/L proteose peptone, 1.5 g/L K_2_HPO_4_, 1.5 g/L MgSO_4_·7H_2_O, and 10 mL/L glycerol) at 28°C for 12 hours (overnight) until reaching an optical density at 600 nm (OD_600_) of 1–2, corresponding to stationary phase. *Bacillus cereus* ATCC 14579, *Pseudomonas aeruginosa, Escherichia coli* K-12 MG1655, and *Klebsiella pneumoniae* CR-HvKP4 were grown in Luria-Bertani (LB) broth at 37°C until achieving an OD_600_ of 1–2 for stationary phase samples. For exponential phase samples, all bacterial strains were first cultured to the stationary phase as described, then subcultured into fresh medium and incubated under the same conditions until reaching an OD_600_ of 0.5–0.6. For *P. syringae* in MM, exponential phase cells were harvested, washed three times with freshly prepared minimal medium(Huynh et al., 1989) (MM), resuspended in MM at an OD_600_ of 0.1–0.2, and cultured for an additional 6 hours.

### baBID-seq library construction

RNA was extracted using RNA isolation Kit V2(Vazyme, #RC112-01). The extracted RNA was processed using DNase I (RNase-free, NEB # M0303S) and collected using RNA Clean & Concentrator™-5 (Zymo #R1014). The RiboRID technique was used for rRNA depletion(Choe et al., 2021). For strains B.cereus and K. pneumoniae, rRNA was depleted with NEBNext® rRNA Depletion Kit (Bacteria) (#E7850L). RNA concentration in each step was tested using Qubit RNA Assays.

One hundred nanograms of RNA generated from the ribosome removal process from each biological replicate were used for the library construction. The fragmentation was optimized to 4 min under 70 °C with the fragmentation buffer used (Invitrogen). For the following steps, we strictly followed the BIDseq protocol(Zhang et al., 2024). The final amplified cDNA are collected and then optimal library fragments are selected using native PAGE gels. 40% Acrylamide/Bis (29:1) 10% native polyacrylamide gel used. The optimal band (175-200bp) used collected. 400 μL of 1x TE buffer and soak the gel thoroughly at 37 °C for 1 h on the thermal shaker at 600 rpm. The gel was crushed and snap-freezed using liquid nitrogen and incubated on a thermal shaker at37 °C at 600rpm for 12 hours. The supernatant collected using Spin-X followed the DNA precipitation. The constructed libraries were sequenced on the Illumina NovaSeq sequencing platform in paired reads mode and single-end reads used for following BID-seq data processing.

pseU-TRACE verification RNA samples (500 ng each for input and bisulfite-treated conditions) were subjected to bisulfite treatment using freshly prepared bisulfite (BS) reagent containing 2.4 M Na_2_SO_3_ and 0.36 M NaHSO_3_, followed by incubation at 70 °C for 3 h. Treated RNA was purified using the Zymo RNA Clean & Concentrator-5 kit. For reverse transcription, 1 μl of 5 μM site-specific RT primer (for either the target Ψ site or the negative control site) was added, and samples were incubated at 65 °C for 5 min and immediately placed on ice. Reverse transcription was then performed using SuperScript IV reverse transcriptase (ThermoFisher #18090050), following the same RT conditions as in the BID-seq protocol. The resulting cDNA was treated with RNase H (NEB #M0297S) at 37 °C for 20 min, then inactivated at 70 °C for 5 min. For splint-ligation, 1 μl of cDNA was mixed with upstream and downstream primers (final concentration 0.01 μM each), and the mixture was annealed using a temperature gradient (90 °C for 1 min, 80 °C for 1 min, 70 °C for 1 min, 60 °C for 1 min, 50 °C for 1 min, and 40 °C for 6 min). SplintR ligase (NEB #M0375S) (2 μl, 1 U) was then added, and ligation was carried out at 40 °C for 60 min, followed by denaturation at 95 °C for 5 min and holding at 12 °C. The reaction was diluted with 40 μl RNase-free H_2_O. Quantitative real-time PCR (qPCR) was performed using a QuantStudio™ 3 PCR System. Each 20 μl reaction contained 2× SYBR Green qPCR Master Mix (MCE #HY-K0501), qPCR forward and reverse primers, diluted ligation product, and RNase/DNase-free water. The qPCR cycling conditions were as follows: 95 °C for 5 min; 40 cycles of 95 °C for 10 s and 60 °C for 35 s; followed by 95 °C for 15 s and 60 °C for 1 min (fluorescence acquisition at a ramp rate of 0.05 °C/s), and a final hold at 4 °C. Ct values were normalized to the negative control site within each replicate, and the normalized treated signal was further normalized to the corresponding input sample to quantify the Ψ modification level. Primer sequence used for detection were listed in **Table 2**.

### Sequencing data processing and analysis

Sequencing data were subjected to a refined bioinformatic workflow adapted from previously established protocols. Raw sequencing reads underwent adapter trimming via Cutadapt (v.3.5) and PCR duplicate elimination using BBMap tools (v.38.73). The filtered reads were initially aligned to a curated repository of non-coding RNA sequences (including rRNAs, tRNAs, and other small RNAs). Subsequently, unmapped reads were subjected to genome alignment with optimized mapping parameters tailored for Ψ detection. The resultant alignment files were processed with Samtools (v.1.13) to generate strand-specific BAM files, which were then interrogated using bam-readcount (v.1.0.1) to quantify nucleotide deletion events and calculate coverage metrics. Ψ modification sites were identified by integrating deletion rate profiles, site coverage, and background signal correction derived from “input” libraries. Bacterial samples from distinct temporal phases were analyzed as discrete entities to minimize batch effects. The quantitative assessment of Ψ levels was achieved by transforming raw deletion ratios according to previously reported calibration curves, yielding high-confidence Ψ fractions at single-nucleotide resolution.

To ensure robust and reliable detection of low-level Ψ sites in mRNA, we applied the following stringent BID-seq filtration criteria to all candidate sites (all sites reported in the **Table.1** passed these thresholds): (1) total sequencing coverage >20 reads in both the bisulfite-treated (BID-seq) libraries (Σdt > 20) and untreated input libraries (Σdi > 20); (2) average deletion number >5 in the treated libraries; (3) average modification fraction >0.02 (2%) in the treated libraries; and (4) average deletion ratio in treated libraries at least two-fold higher than in input libraries. We consider sites with stoichiometry thresholds > 0.5 as highly modified sites.

### TPM calculation

The TPM value for a specific transcript *i* is calculated as:

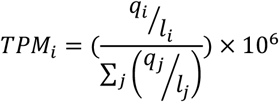

*q*_*i*_ was the number of reads mapped to transcript i. *l*_*i*_ is the length of transcript I (in nucleotides). 10^6^ was the scaling factor to express the result as transcripts per million(Zhao et al., 2021).

### Evolutionary analysis

Orthologous gene analysis was performed using OrthoFinder(Emms and Kelly, 2019) software to identify homologous gene clusters across the five bacterial strains. The analysis incorporated complete genome sequences and their corresponding GFF (General Feature Format) annotation files with default parameters. The resulting orthologous gene clusters were subsequently utilized for KEGG (Kyoto Encyclopedia of Genes and Genomes) pathway analysis. Genes that were not clustered in the orthology analysis were excluded from the KEGG pathway mapping to ensure reliable results. Functional annotations were performed using eggNOG-mapper v2(Cantalapiedra et al., 2021).

### RNA structure analysis

Pseudouridine modification sites were filtered with a fraction of larger than 2%. RNA sequences were generated by extracting 20nt upstream and downstream of each modification site, yielding 41 nucleotide sequences. These sequences were subsequently used for RNA secondary structure prediction with MXfold2(Sato et al., 2021) and sequence clustering analysis with RNAclust(Engelhardt et al., 2010). Results analysis and visualization were performed using customized scripts written in R.

### pseU_GNN

#### Training data

The negative control dataset was constructed by scanning bacterial genomes for pseudouridylation-compatible sequence motifs absent from the experimentally verified modification sites in baBID-seq data, then generating sequence segments of 41, 61, and 81 nucleotides with these unmodified motifs at their centres. MXfold2(Sato et al., 2021) is used to predict the secondary structure of all sequences as input for training and validation process. 3,377 high-confidence Ψ sites (with fraction values >2%) as positive samples. The negative samples consisted of 3,400 randomly selected sites that contained the unique Ψ motif but showed no evidence of Ψ deposition in our experimental data. The dataset was divided into 4,744 training samples, 1,016 test samples, and 1,017 validation samples for each sequence length.

### Model Architecture

The model architecture comprises three main components. First, two dense graph encoders were used to extract RNA secondary structure features derived from MXfold2 predictions and from one-hot–encoded RNA sequences, respectively. Second, the one-hot–encoded sequence was further processed by a convolutional layer for local feature extraction, followed by two bidirectional LSTM layers to capture long-range dependencies. Positional encoding was then applied before forwarding the representations to a single-layer multi-head Transformer block.

Finally, features extracted from the structure-based and sequence-based modules were integrated via weighted concatenation and passed through a residual block consisting of fully connected layers. A sigmoid activation function was applied to the final output to predict the probability of pseudouridine modification.

### Training and Evaluation

The pseU_NN was trained using binary cross-entropy loss with the Adam optimizer. We implemented a dynamic learning rate scheduler that adjusted the rate based on validation AUC-ROC performance.

For the final evaluation, we utilized an independent test dataset, with F1 scores and precision-recall curves guiding prediction threshold selection. This balanced approach ensured optimal sensitivity and specificity - critical in biological classification systems where both false positives and false negatives carry significant consequences.

The F1 score is the harmonic mean of precision and recall. The definition of true positive (TP), true negative (TN), false positives (FP) and false negative (FN) with shifting was adopted in previous studies(Wang et al., 2023; Yu et al., 2021).

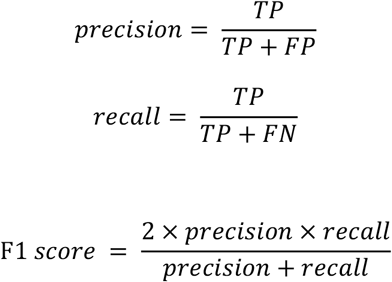

The accuracy calculated using standard definitions:

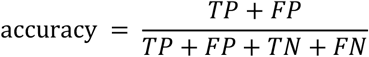

The Matthews correlation coefficient (MCC) is calculated using the following formula:

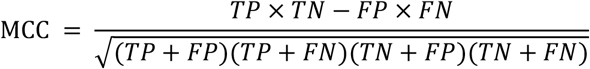

## Acknowledgments

This study was funded by the Guangdong Major Project of Basic and Applied Basic Research (2020B0301030005), National Natural Science Foundation of China (32172358), General Research Funds of Hong Kong (21103018, 11101619, and 11102720) and Early Career Scheme of Hong Kong (26103623). The funders had no role in study design, data collection, interpretation, or the decision to submit the work for publication.

## Data availability

All sequencing data files have been submitted to the National Center for Biotechnology Information (NCBI) Gene Expression Omnibus (GEO) database with the reference code of GSE292335.

## Code availability

The code for pseU_NN used for this paper are available at https://github.com/Dylan-LT/pseU_NN.git

## Author contributions

X.D. and L.X. conceived the original idea. X.D. and L.-S.Z. supervised the project. L.X. and Y.Z. designed and performed experiments. L.X., S.S., and Y.S. performed bioinformatic analysis and prepared figures. L.X. and Z.G. designed pseU_NN. B.L., Z.G., and R.L. assisted with experiments. L.X., L.-S.Z., and X.D. wrote the manuscript. All the authors read the paper and agreed with the final version.

---

**Supplementary Figure 1.**
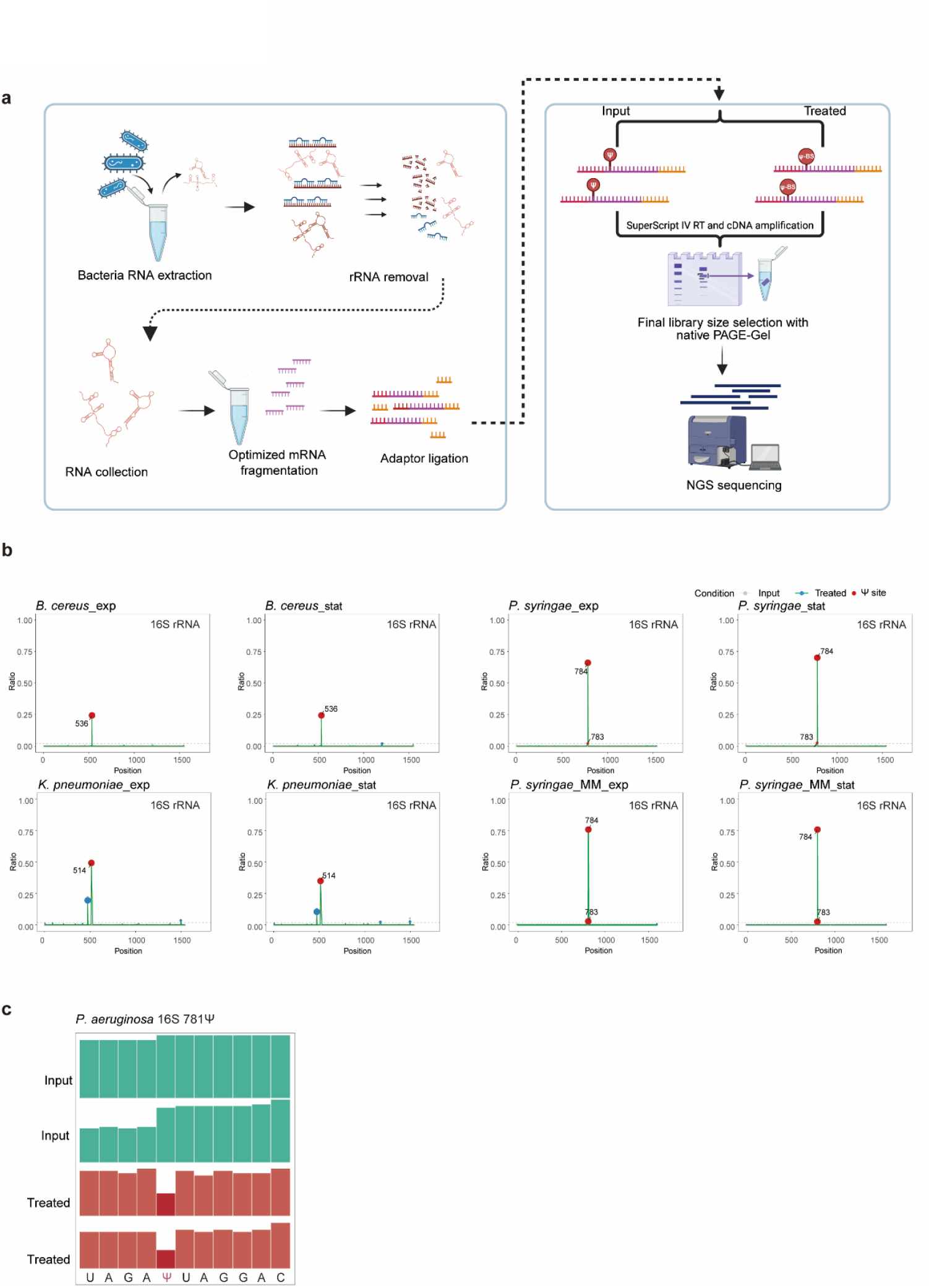
baBID-seq workflow and detection of Ψ sites on 16S rRNA. **a** Schematic overview of the baBID-seq workflow optimized for bacterial RNA, adapted from the original BID-seq protocol. **b** Distribution of detected Ψ sites on 16S rRNA across different bacterial strains, with the y-axis showing the deletion ratios in input and bisulfite-treated samples. **c** Integrative Genomics Viewer (IGV) snapshot illustrating the detected Ψ781 site in *P. aeruginosa* 16S rRNA.

**Supplementary Figure 2.**
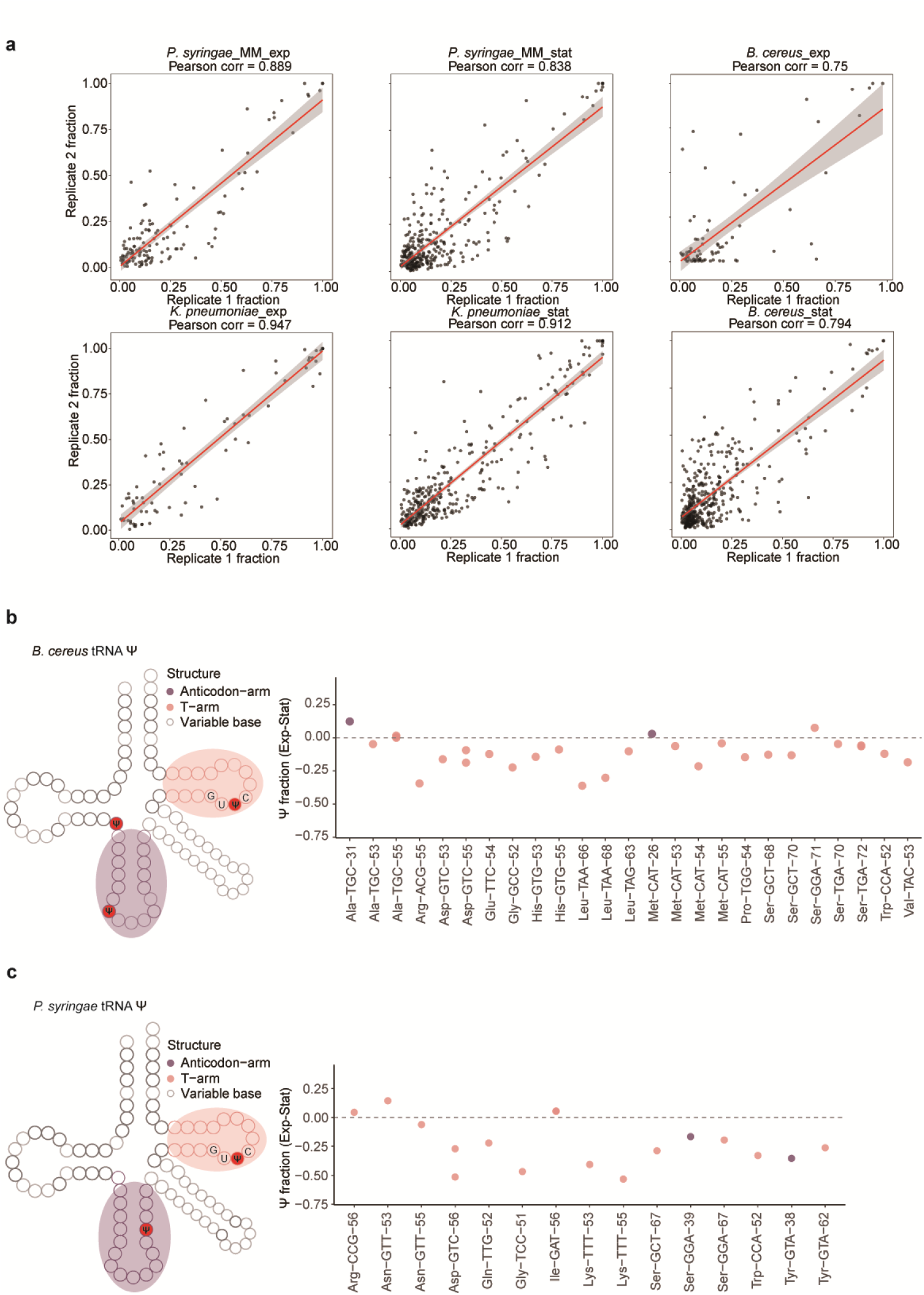
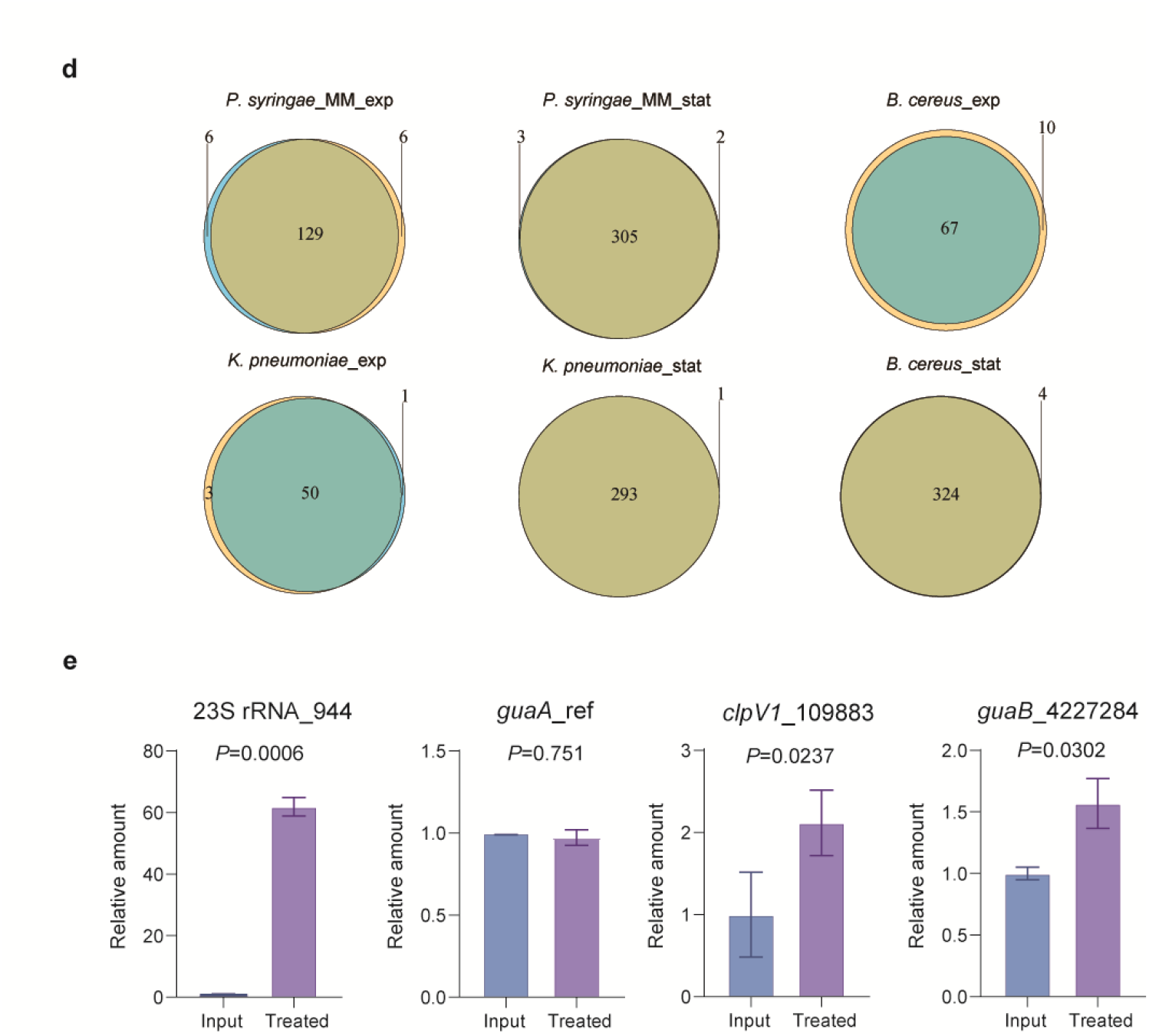
Reproducibility of Ψ detection and dynamic Ψ modification in tRNA. **a** Pearson correlation of Ψ modification fractions between two independent biological replicates for each strain. b**-c** Changes in tRNA Ψ fractions in *B. cereus* (**c**) and *P. syringae* (**d**) between exponential and stationary growth phases. The y-axis represents the Ψ fraction difference (exponential phase minus stationary phase). **d** Overlap of identified mRNA Ψ sites between biological replicates for each strain. **e** Experimental validation of Ψ sites in *P. aeruginosa* PAO1 using the pseU-TRACE method. From right to left, the panels show validation of the Ψ site at position 944 on 23S rRNA, a negative control site from the *guaA* gene, a Ψ site in *clpV1* (chromosomal position 109,883), and a Ψ site located in the intergenic region between *guaA* and *guaB* (chromosomal position 4,227,284).

**Supplementary Figure 3.**
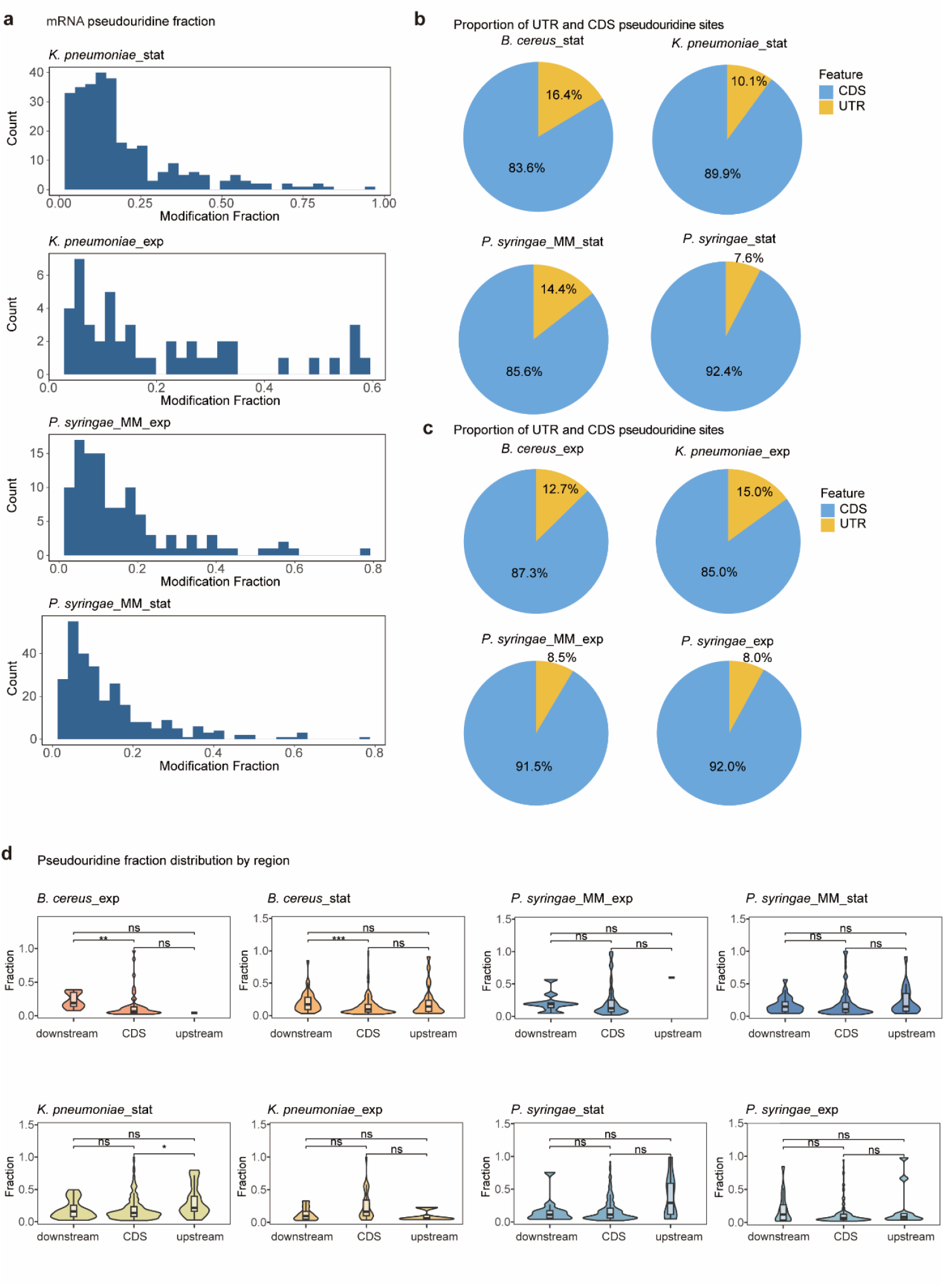
Distribution and fraction of mRNA Ψ modifications across transcript regions. **a** Distribution of mRNA Ψ fraction values in *K. pneumoniae* and *P. syringae* under minimal medium (MM) conditions. **b** Distribution of Ψ modifications between untranslated regions (UTRs) and coding sequences (CDSs) during the stationary growth phase and (**c**) exponential growth phase. **d** Distribution of Ψ fraction values in upstream and downstream UTRs of mRNAs across exponential and stationary growth phases.

**Supplementary Figure 4.**
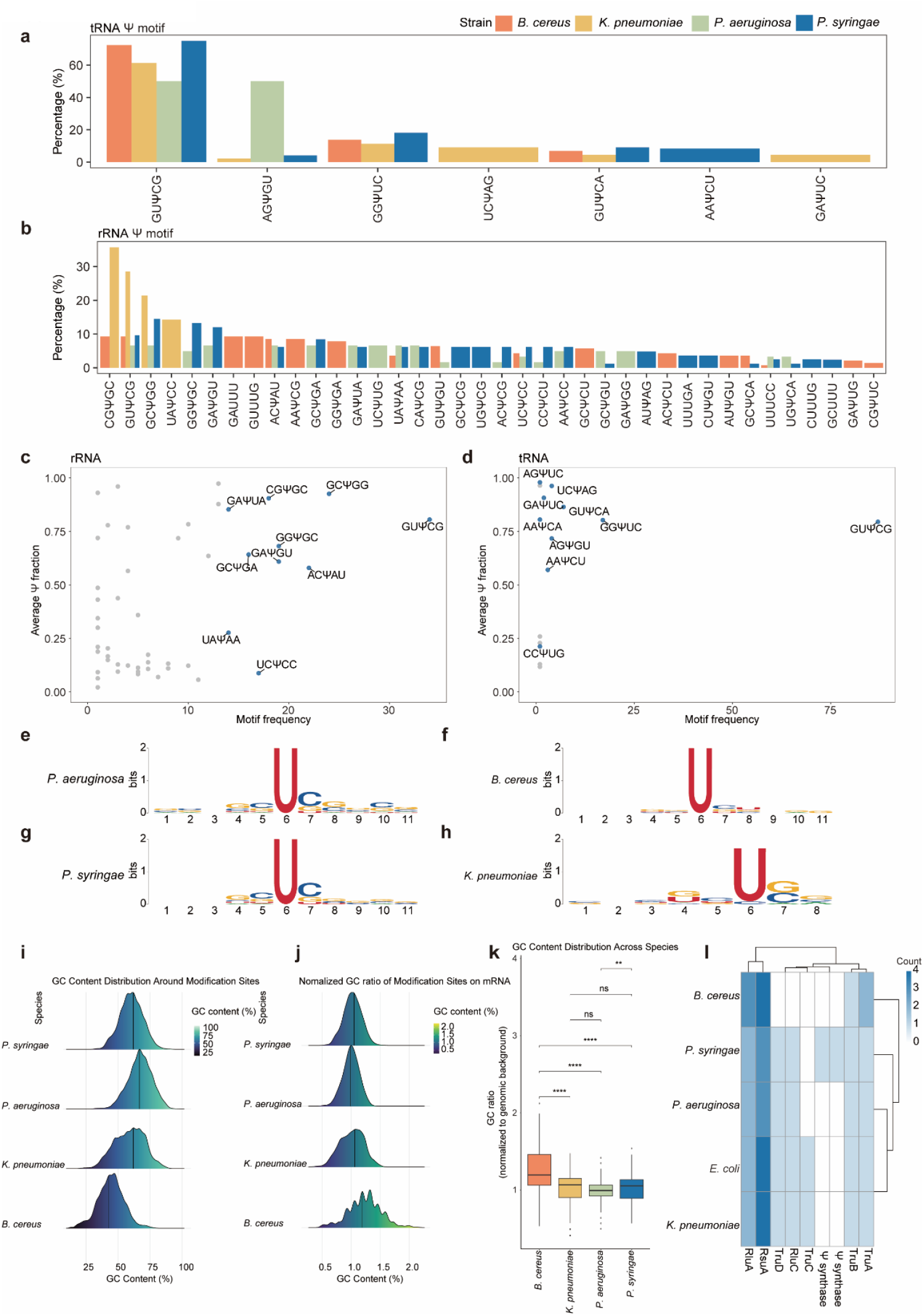
Motif pattern of Ψ modification. **a** The Ψ motif ratio in tRNA across four strains. Motif ratios are calculated by dividing the count of each specific 5-mer motif centered on Ψ by the total number of motifs detected in each individual strain tRNA. **b** The Ψ motif distribution ratio in ribosomal RNA (rRNA) across four bacterial strains. Motif ratios are calculated by dividing the count of each specific 5-mer motif centered on Ψ by the total number of motifs detected in each individual strain rRNA. **c** Scatter plot displaying the average Ψ fraction values for different sequence motifs in ribosomal RNA. Each point represents a specific sequence motif, with its position on the y-axis indicating the average Ψ fraction value across all instances of that motif in rRNA. **d** Similar to **c**, each point represents a specific motif, with its position on the y-axis indicating the average Ψ fraction value across all instances of that motif in tRNA. **e-h** Pseudouridylation motifs identified on mRNA transcripts from four bacterial strains using MEME. **i** Density plot displaying GC composition within 10-base pairs flanking Ψ sites in mRNA. **j** GC composition within 10-base pairs flanking Ψ sites in mRNA normalized to the background GC ratio. **k** Box plot comparing the GC ratio (normalized to genome background) around Ψ sites across different bacterial strains. **l** Hierarchical clustering plot showing gene orthology analysis of all pseudouridine synthases across bacterial strains. The colour intensity represents clustered gene counts across genomes.

**Supplementary Figure 5.**
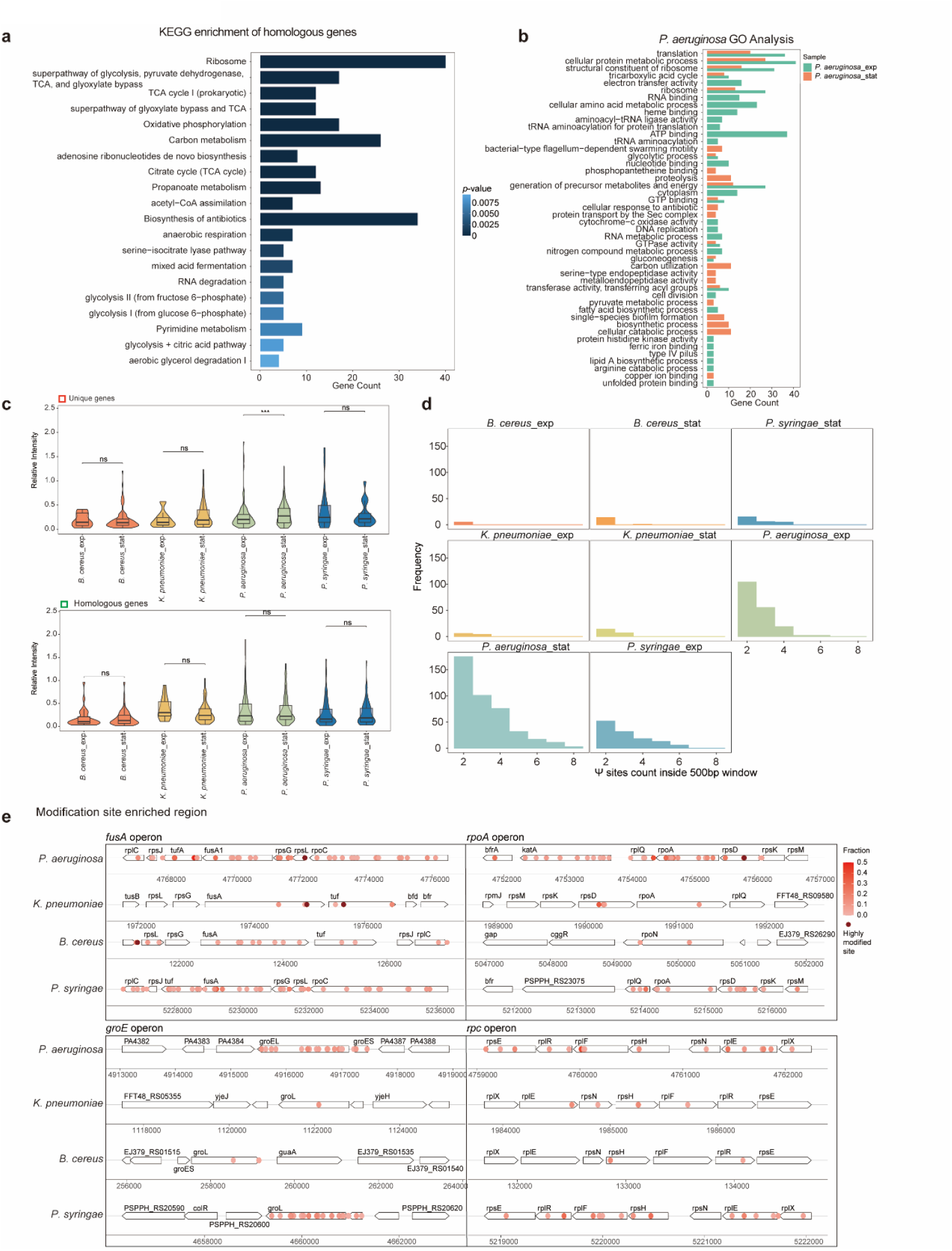
Comprehensive analysis showing functional enrichment of Ψ-containing transcripts and Ψ enriched region. **a** The bar plot illustrates the top 20 enriched KEGG pathways, ranked by *p*-value, derived from Ψ modified homologous genes identified in at least two strains. **b** The Gene Ontology analysis of Ψ-mRNA in exponential and stationary phases of *P. aeruginosa*. **c** Comparison of Ψ intensity patterns between strain-specific and homologous gene sets during two different growth phases. **d** Analysis of pseudouridylation density showing the number of Ψ sites occurring within 500nt windows centered on each modified position. **e** Ψ modification-enriched regions detected across bacterial strains. The x-axis represents positions on the bacterial chromosome. Dark red indicates highly modified Ψ sites, defined as positions where the Ψ fraction exceeds 50%.

**Supplementary Figure 6.**
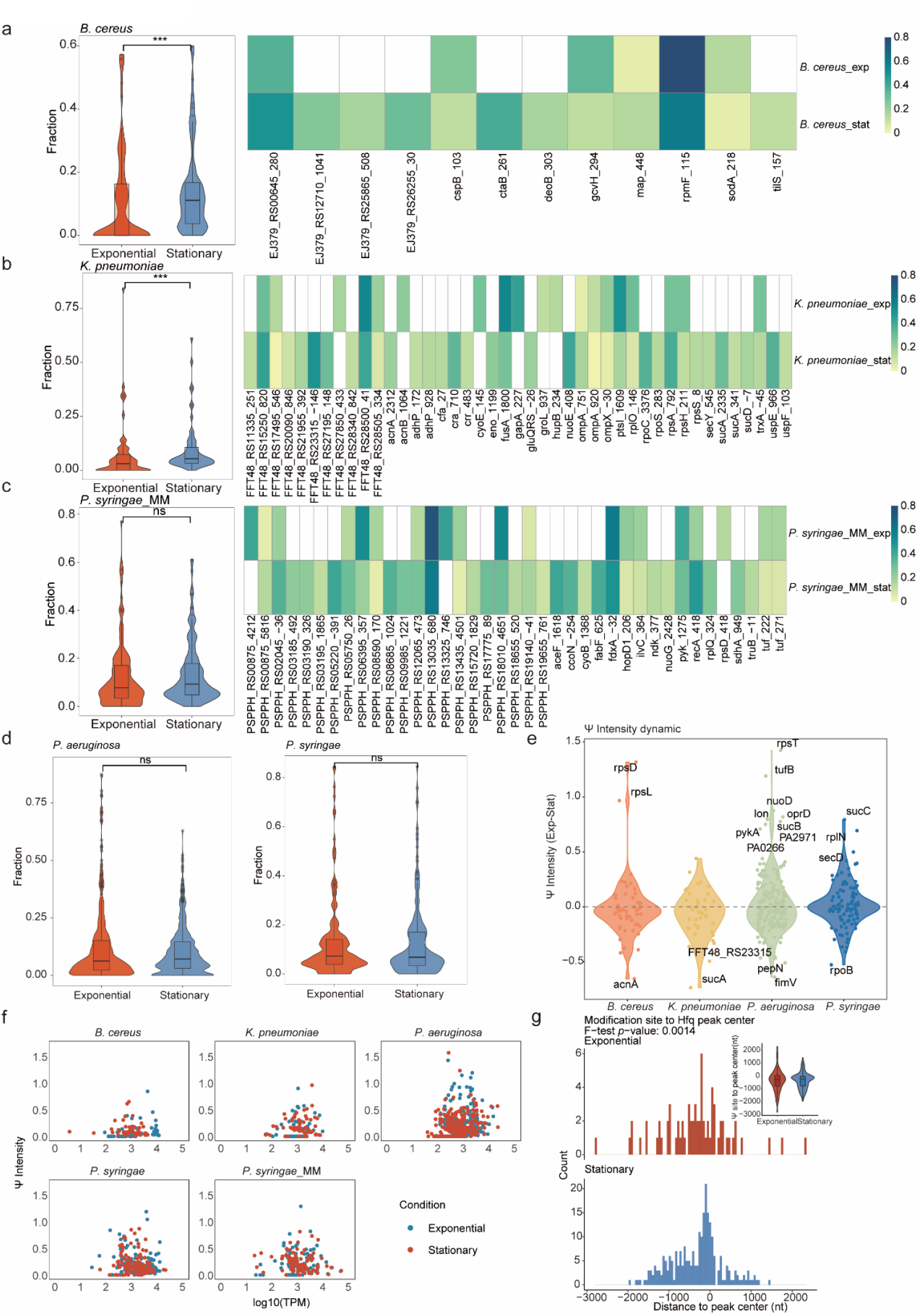
Growth state-dependent dynamics of Ψ modification fraction. **a-c** Paired visualization showing Ψ dynamics across growth phases, with a violin plot depicting the distribution of Ψ fractions in mRNAs during two distinct growth phases (left), and a corresponding heatmap displaying the specific Ψ modification sites dynamic detected across bacterial strains at these same growth phases (right). **d** The Ψ fraction distribution on mRNA in *P. aeruginosa* (left) and *P. syringae* (right) at two growth phases. **e** Violin plot quantifying the differential pseudouridylation intensity (exponential phase Ψ intensity minus stationary phase Ψ intensity) of specific genes at two growth phases. **f** Distribution plot illustrating the spatial relationship between Ψ modification sites and Hfq binding peak centres during two distinct growth phases. The violin plot reveals that the distribution of minimum distances between Ψ sites and Hfq binding centres varies with statistical significance (*p-*value = 0.0014) between growth phases, as determined by an F-test. **g** The scatter plot depicts the relationship between Ψ modification intensity and gene expression levels (log10(TPM)) under two distinct growth conditions.

**Supplementary Figure 7.**
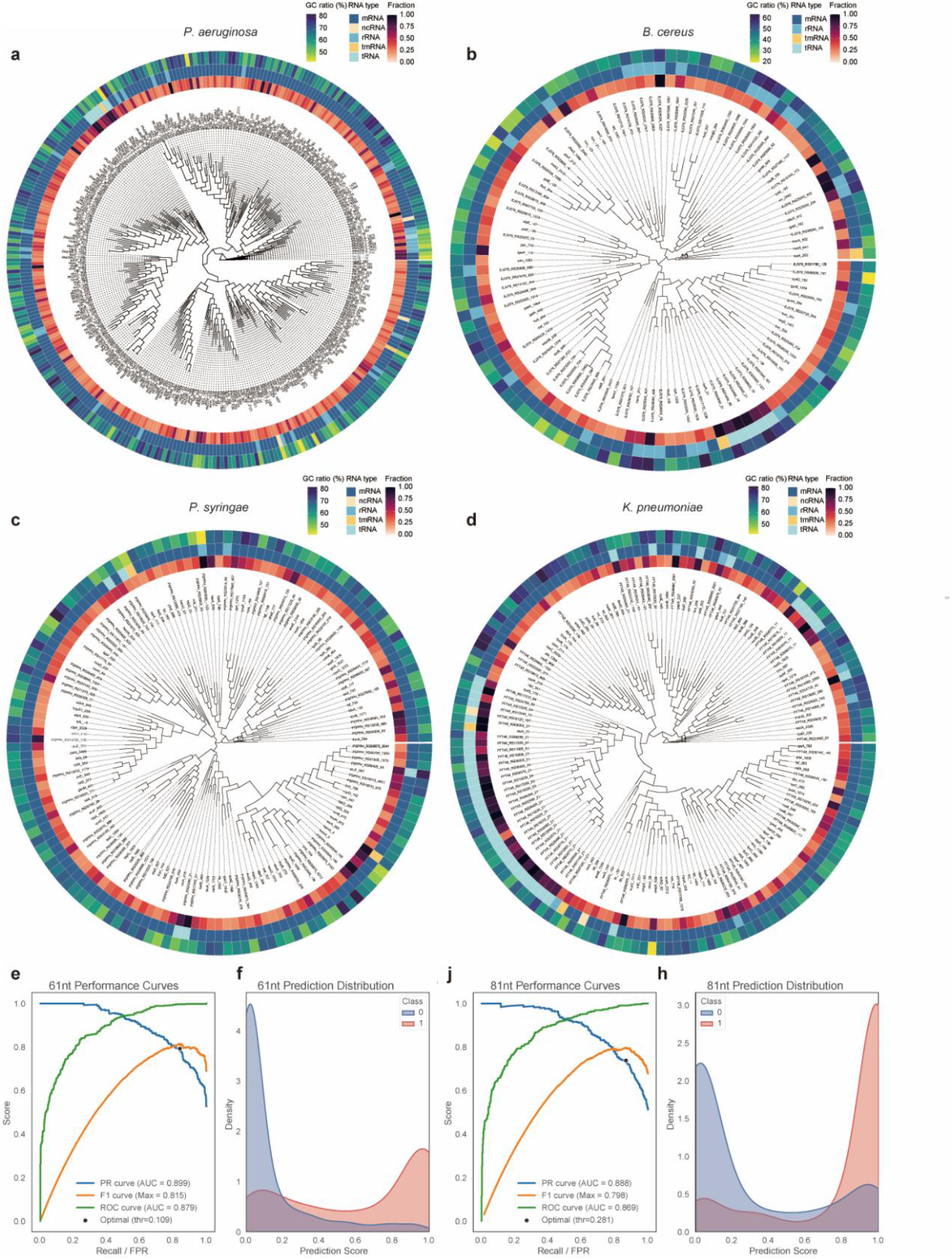
The structure and sequence clustering analysis of RNA molecules containing Ψ modifications. **a-d** Sequence and structure clustering of 41-nucleotide RNA segments centered by Ψ modification sites with fraction values greater than 0.1. The circular visualization comprises three concentric layers: the inner layer displays the Ψ fraction value, the middle layer designates RNA type, and the outer layer represents the GC ratio of each 41-nucleotide RNA segment. **e** Multi-metric assessment showing the precision– recall curve (AUC = 0.899), F1 score curve, and ROC curve (AUC = 0.879) of the pseU_NN model on the 61-nt validation dataset. The right panel **f** shows the distribution of pseU_NN prediction scores on the 61-nt validation dataset. **j**,**h** Similar to panels **e**,**f**, multi-metric assessment showing the precision–recall curve (AUC = 0.888), F1 score curve, and ROC curve (AUC = 0.869) of the pseU_NN model on the 81-nt validation dataset, and the right panel shows the distribution of pseU_NN prediction scores on the 81-nt validation dataset.

